# Naïve and *in vitro*-activated primary mouse CD8^+^ T cells retain *in vivo* immune responsiveness after electroporation-based CRISPR/Cas9 genetic engineering

**DOI:** 10.1101/2021.09.14.460345

**Authors:** Petra Pfenninger, Laura Yerly, Jun Abe

## Abstract

CRISPR/Cas9 technology has revolutionized genetic engineering of primary cells. Although its use is gaining momentum in studies on CD8^+^ T cell biology, it remains elusive to what extent CRISPR/Cas9 affects *in vivo* function of CD8^+^ T cells. Here, we optimized nucleofection-based CRISPR/Cas9 genetic engineering of naïve and *in vitro*-activated primary mouse CD8^+^ T cells and tested their *in vivo* immune responses. Nucleofection of naïve CD8^+^ T cells preserved their *in vivo* antiviral immune responsiveness to an extent that is indistinguishable from non-nucleofected cells, whereas *in vitro* activation of CD8^+^ T cells prior to nucleofection led to slightly impaired expansion/survival. Of note, different target proteins displayed distinct decay rates after gene editing. This is in stark contrast to a comparable period of time required to complete gene inactivation. Thus, for optimal experimental design, it is crucial to determine the kinetics of the loss of target gene product to adapt incubation period after gene editing. In sum, nucleofection-based CRISPR/Cas9 genome editing achieves efficient and rapid generation of mutant CD8^+^ T cells without imposing detrimental constraints on their *in vivo* functions.

## Introduction

Mutant mouse lines have played essential roles in immunology to identify the function of many genes in the immune system. Although such mutant lines continue to be powerful and highly useful to date, the generation of desirable mouse lines often requires time-consuming interbreeding of multiple mouse lines. To study T cell biology *in vivo* using adoptive transfer, targeted mutation alone is not sufficient to track the antigen-specific response of transferred T cells in the host for a long period of time; mutant T cells also have to carry a congenic marker and a transgenic T cell antigen receptor (TCR) specific for a model antigen. When the gene of interest interferes with T cell development, there is an additional need for its inducible expression or deletion. In such cases, mutant mouse lines have to carry yet another locus for Cre recombinase or other mechanisms that allow for inducible modification of the genome. Thus, it is common to introduce 3–4 congenic or mutant loci to study the function of a gene in T cells.

The advent of CRISPR/Cas9-based genetic engineering technology in the last decade has revolutionized this situation (1–3). While the faster generation of mutant mouse lines is a great advantage of CRISPR/Cas9 over previous techniques, its use further extends to the direct generation of mutant cells. When combined with TCR-transgenic, congenic mice as the source of T cells, CRISPR/Cas9 technology permits to skip the time-consuming interbreeding before performing adoptive transfer experiments. Until recently, however, genetic engineering of primary T cells relied almost completely on gene transduction using viral vectors, because it is notoriously difficult to introduce exogenous genetic elements into non-dividing primary T cells (4). One of the drawbacks of this approach is that T cell activation is a prerequisite for the integration of transgene(s) into the genome (4, 5), hampering its use to study functions of the gene of interest in naïve T cells or during T cell activation. Furthermore, sustained expression of guide RNA (gRNA) and Cas9 nuclease in transduced cells can aggravate off-target effects (6–8) and potentially lead to rejection of cells after adoptive transfer.

To overcome the challenges associated with viral transduction, several recent studies deployed Nucleofector™ technology to deliver pre-assembled ribonucleoprotein (RNP) complex, comprising gRNA and recombinant Cas9, into T cells irrespective of their cell cycle stage (4, 9, 10). This has allowed to generate knockout cells within approximately a week without prior activation of T cells. One of these reports aimed to optimize nucleofection conditions for naïve and *in vitro*-activated primary CD4^+^ and CD8^+^ T cells (4). Yet, it remains elusive whether different nucleofection conditions affect *in vivo* immune responsiveness of CRISPR/Cas9-engineered T cells.

In this study, we re-evaluated optimal nucleofection conditions for naïve and *in vitro*-activated primary mouse CD8^+^ T cells with a revised screening panel based on a previous study (4) and inputs from the manufacturer. Then, we tested their immune responsiveness after they undergo all the processes of nucleofection-based CRISPR/Cas9 genetic engineering (pre-stimulation, nucleofection and post-nucleofection incubation). Furthermore, we tested whether use of a second target as a reporter allows for enrichment of cells in which the primary target locus is successfully edited, as a way to isolate desirably engineered cells.

## Methods

### Animals

B6.SJL-Ptprc^a^ (CD45.1; MGI 2164701) (11), C57BL/6-Tg(TcraTcrb)1100Mjb/J (OT-I; MGI 3054907) (12), Tg(UBC-GFP)30Scha (Ubc-GFP; MGI 305178) (13), Tg(Zp3-cre)82Knw (Zp3-Cre; MGI 4888024) (14), Gt(ROSA)26Sor^tm14(CAG-tdTomato)Hze^ (Ai14; MGI 3809524) (15) and Dock2^tm1Ysfk^ (DOCK2-GFP; MGI 3772945) (16) mice were bred under specific pathogen-free conditions at the University of Fribourg. To obtain mice expressing tandem-dimer Tomato (tdT) driven by CAG promoter (tdT mice), Ai14 mice were crossed with Zp3-Cre mice to excise the floxed stop codon in the Ai14 alleles. Resultant tdT-expressing mice were backcrossed with wild-type C57BL/6J (B6J) mice to remove Zp3-Cre transgene. GFP-, tdT- or CD45.1-expressing OT-I mice were derived by crossing OT-I mice with Ubc-GFP, tdT or CD45.1 mice, respectively. Female and male B6J mice at the age of 4–6 weeks old were purchased from Janvier (Le Genest-Saint-Isle, France) and used for screening and as recipient of sex-matched OT-I cells. All animal experimentations have been approved by the Cantonal Committees for Animal Experimentation and performed in accordance with the federal guidelines.

### T cell isolation

Single cell suspension of spleen and peripheral lymph nodes (LNs) was prepared by mincing them using a 70-µm cell strainer. Untouched CD8^+^ T cells were magnetically isolated using EasySep™ Mouse CD8^+^ T Cell Isolation Kit (STEMCELL Technologies, Grenoble, France) or MojoSort™ Mouse CD8 T Cell Isolation Kit (BioLegend, San Diego, CA). Purity of the isolate CD8^+^ cells constantly exceeded 95%.

### T cell stimulation before nucleofection

For activated CD8^+^ T cells, T-25 or T-75 flasks were coated with 5 µg/mL anti-CD3ε antibodies (clone 145-2C11; 100340, BioLegend) in phosphate-buffered saline (PBS) for 16–24 hr at 4°C before seeding CD8^+^ T cells. Immediately after magnetic isolation, CD8^+^ T cells were resuspended in complete medium (RPMI1640 medium supplemented with 10% fetal calf serum (FCS), 2 mM L-glutamine, 0.1 mM non-essential amino acids, 1 mM sodium pyruvate, 100 U/mL penicillin, 0.1 mg/mL streptomycin, 50 µM 2-mercaptoethanol and 10 mM HEPES) and seeded in the antibody-coated flask at 3–5×10^5^ cells/cm^2^ together with soluble 1 µg/mL anti-CD28 antibodies (clone 37.51; 102116, BioLegend). Cells were cultured for 48 hr at 37°C in a humidified 5% CO_2_ atmosphere. For naïve CD8^+^ T cells, magnetically isolated T cells were resuspended in the complete medium and were seeded in a 10-cm petri dish or T-75 flask at 1–2×10^6^ cells/cm^2^ and cultured in the presence of 20 ng/mL recombinant mouse interleukin-7 (rmIL-7; 402-ML-020/CF, R&D Systems, Minneapolis, MN) for 24 hr at 37°C in a humidified 5% CO_2_ atmosphere.

### Nucleofection

CD90 crisprRNAs (crRNA) were designed in the previous study (4). DOCK2 crRNA were designed using DESKGEN online tool (www.deskgen.com; discontinued). Alt-R^®^ CRISPR-Cas9 crRNA (custom design) and trans-activator RNA (tracrRNA) (1072534) were purchased from Integrated DNA Technologies (Coralville, IA) and reconstituted at 100 µM with nuclease free duplex buffer (Integrated DNA Technologies). Nucleofection was performed as described previously (4) with some modifications. In brief, one microliter each of crRNA and tracrRNA were annealed to form gRNA at 95°C for 10 min using a thermal cycler and cooled to room temperature. Annealed gRNA was mixed with TrueCut Cas9 v2 (A36499, Thermo Fisher Scientific, Basel, Switzerland) at a ratio of gRNA : Cas9 = 1.8 µL : 1.2 µL (equivalent to 90 pmol : 36 pmol) and left at room temperature for > 10 min to generate RNP complex. Pre-stimulated T cells were harvested, counted, and spun down at 80×***g*** for 7 min. Pellets were resuspended in Primary Cell 4D-Nucleofector™ X Kit S (Lonza) buffer solution at a cell concentration of 1–4×10^6^ cells in 20 µL. The entire cell suspension was mixed with 3 µL per complex RNP solution and added to Nucleocuvette™ strip well. Cells were then nucleofected using a 4D-Nucleofector™ with X-Unit (V4XP-4032 and V4XP-9096, Lonza, Basel, Switzerland). Up to three RNP complexes in 9 µL were used per reaction. Immediately after nucleofection, one hundred microliter of pre-warmed complete medium containing 10 ng/mL recombinant mouse IL-2 (rmIL-2; 402-ML-020/CF, R&D Systems) or 20 ng/mL rmIL-7 was added to each Nucleocuvette™ strip well for activated or naïve CD8^+^ T cells, respectively. Then, cells were gently mixed by pipetting and aliquoted into a flat-bottom 96-well plate. Cells were cultured in a total volume of 250 µL complete medium containing rmIL-2 or rmIL-7 at 1×10^5^ and 2×10^6^ cells per well for activated and naïve CD8^+^ T cells, respectively, for 2–10 days at 37°C in a humidified 5% CO_2_ atmosphere. crRNAs used in this study are listed in **Table 1**. Buffer and pulse code are indicated in the corresponding figure legends.

**Table 1.**
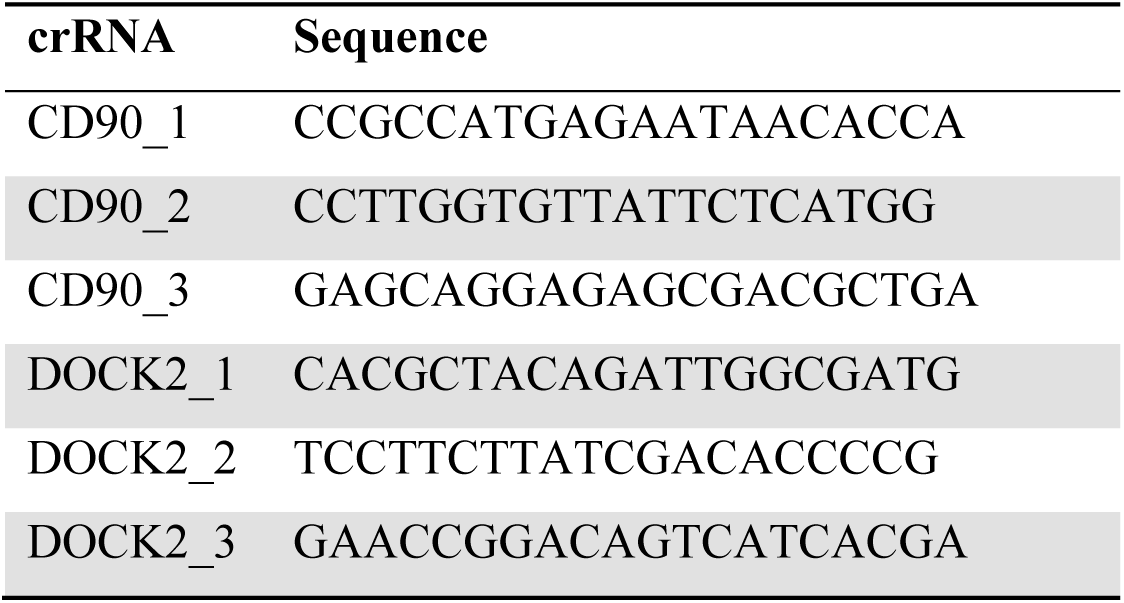
Sequence of crRNAs

### Viral infection

For activated CD8^+^ T cells, mice subcutaneously infected with 5×10^4^ plaque-forming unit (pfu) herpes simplex virus-1 expressing tdT and ovalbumin (HSV-OVA) (17) via hock (18) received a mix of 1.5×10^5^ each of nucleofected or control *in vitro*-activated OT-I cells on day 4 post-infection. For naïve CD8^+^ T cells, mice were intravenously injected with a total of 1.5×10^4^ nucleofected OT-I cells comprising 5×10^3^ each of nucleofected and control OT-I cell populations. One day later, mice were subcutaneously infected with 5×10^4^ pfu HSV-OVA in 20 µL PBS via hock. Each mouse received three OT-I cell populations that were differently nucleofected (or left non-nucleofected) and marked by the expression of GFP, tdT or CD45.1.

### Antibodies

Following antibodies were purchased from BioLegend and used for staining: Brilliant Violet 421™ (BV421)-conjugated anti-CD45.1 (clone A20; 110732) and anti-TNF-α (clone MP6-XT22; 506328), Brilliant Violet 605™ (BV605)-conjugated anti-CD44 (clone IM7; 103047) and anti-CXCR3 (clone CXCR3-173; 126523), Brilliant Violet 711™ (BV711)-conjugated anti-CD45.1 (clone A20; 110739) and anti-IFN-γ (clone XMG1.2; 505836), PerCP/Cy5.5-conjugated anti-CD43 activation associated glycoform (clone 1B11; 121223) and anti-CD90.2 (clone 53-2.1; 105337), PE/Cy7-conjugated anti-CD45.1 (clone A20; 110729), anti-CD90.2 (clone 53-2.1; 140309) and anti-KLRG1 (clone 2F1/KLRG1; 138415) and anti-CD127 (clone A7R34; 135012), Alexa Fluor™ 700 (AF700)-conjugated anti-CD90.2 (clone 53-2.1; 140323), APC/Fire™ 750-conjugated anti-CD8α (clone 53-6.7; 100766), and Ultra-LEAF™ purified anti-CD16/32 (clone 93; 101330). APC-conjugated anti-IL-2 (clone JES6-5H4; 554429) was purchased from BD Biosciences. Unlabeled anti-CD16/32 (clone 2.4G2; 60161) was purchased from STEMCELL Technologies. Data were acquired using an LSRFortessa (BD Biosciences) and analyzed using FlowJo™ software (FlowJo, Ashland, OR).

### Flow cytometry

For samples from virus-infected mice, popliteal LNs (popLN) and spleen were harvested and minced using a 70-µm cell strainer. Single cell suspension of peripheral blood leukocytes (PBL) was prepared from whole blood by lysing red blood cells. Cells were counted and plated at ≤ 3×10^6^ cells per well in a round-bottom 96 well plate for antibody staining. First, cells were resuspended in 15 µL of 2 µg/mL anti-CD16/32 in staining buffer (PBS containing 2% FCS, 2 mM EDTA and 0.05% (w/v) NaN_3_) and incubated for 5 min on ice to block Fc receptors. Fifteen microliters of antibody mix prepared at twice higher concentration than the working concentration was added to cells without washing away anti-CD16/32 and incubated for 30 min on ice. After staining, cells were washed twice with staining buffer and resuspended in the staining buffer containing 0.5 µg/mL propidium iodide for dead cell exclusion. For cells derived from virus-infected mice, Zombie Red™ Fixable Viability Dye (423110, BioLegend) was added to anti-CD16/32 solution at a 1 : 100 dilution for dead cell exclusion and fixed with 4% paraformaldehyde (15710, Electron Microscopy Sciences, Lucerne, Switzerland) in PBS for 20 min on ice after staining.

### Intracellular cytokine staining

For intracellular cytokine staining, cells were restimulated with 1 µM SIINFEKL (BAP-201, EMC Microcollections, Tübingen, Germany) and 5 µg/mL brefeldin A (B6542, Sigma-Aldrich, St. Louis, MO) in 250 µL complete medium for 5 hr at 37°C in a humidified 5% CO_2_ atmosphere before staining. After surface staining, dead cell labeling with Zombie Red™ dye and fixation, cells were permeabilized by resuspending in 200 µL PermWash buffer (554723, BD Biosciences, San Jose, CA) and labeled with antibodies against cytokines diluted in PermWash buffer.

### Cell sorting

DOCK2-GFP CD8^+^ T cells were nucleofected with three CD90 crRNAs with or without two DOCK2 crRNAs (**Table 1**) on day 0. On day 7 or 10 after nucleofection, cells were harvested and stained with APC/Fire™ 750-conjugated anti-CD8α and PE/Cy7-conjugated anti-CD90.2. After two rounds of wash, cells were resuspended in staining buffer containing 0.5 µg/mL propidium iodide and sorted into CD90^lo^ and CD90^hi^ fractions using a FACSAria™ Fusion cell sorter (BD Biosciences). Sorted cells were collected in the complete medium and processed for total RNA isolation immediately after sorting.

### Quantitative PCR

Total RNA isolation and cDNA preparation from sorted cells were performed using TRI Reagent™ (12044977, Sigma-Aldrich) and PrimeScript™ RT Reagent Kit (RR037B, Takara Bio, Saint-Germaine-en-Laye, France), respectively, according to the manufacturer’s instruction. cDNA was prepared using 200 ng total RNA and random primers. Quantitative PCR was performed in duplicates with 20 ng template cDNA per reaction using KAPA SYBR^®^ FAST (KK4603, Roche, Basel, Switzerland) and a CFX96™ Real Time System C1000 Touch™ Thermal Cycler (BioRad, Hercules, CA). Cycling condition was 95°C for 5 min, followed by 40 cycles of 95°C for 15 sec and 60°C for 60 sec. After the completion of PCR reaction, amplification specificity was confirmed by generating dissociation curves. Relative expression values were calculated by ΔΔCT method. First, relative expression levels of *Dock2* were calculated as 2^−ΔCt^, where ΔCt value represents the difference in Ct values between *Dock2* and *Gapdh* (Ct(*Dock2*) – Ct(*Gapdh*)). Then, *Dock2* expression levels were normalized to the level in control, DOCK2 non-targeted samples. Following primers were purchased from Microsynth (St. Gallen, Switzerland): *Dock2*-Fwd, 5’-GGCTCATAGGATTCTCCATCCG-3’; *Dock2*-Rev, 5’-GATTGGGATGGTGGCTTTCCTG-3’; *Gapdh-*Fwd, 5’-AGAACATCATCCCTGCATCC-3’; *Gapdh*-Rev, 5’-TCATCATACTTGGCAGGTTTCTC-3’.

### Statistics

Graph preparation, statistical tests and calculation of Pearson correlation coefficient were performed using GraphPad Prism software (GraphPad Software; San Diego, CA). Thick lines in scatter plots and error bars represent mean and standard deviation, respectively. Differences among two or more groups were considered statistically significant when *p* < 0.05. Statistical tests used for comparison are indicated in the corresponding figure legends.

## Results

### CRISPR/Cas9-mediated genetic engineering of *in vitro*-activated primary mouse CD8^+^ T cells

First, we examined the optimal nucleofection conditions for *in vitro*-activated mouse CD8^+^ T cells. Here, we chose CD90 as the model target because of its uniform expression on the surface of CD8^+^ T cells, which allows for its easy detection by flow cytometry. For nucleofection, we selected seven electric pulse codes based on a previous report by Seki and Rutz (4) and recommendation from the manufacturer, Lonza: CA137, CM137, DN100, DN107, DO100, DS138 and DV100. Seki and Rutz used pulse codes indicated in the manufacturer’s instruction for the optimization kit for primary cells. Because this recommendation is not tailored for mouse T cells, we surmised that the previous screening panel might have missed certain pulse codes suitable for mouse CD8^+^ T cells. Therefore, we partly revised the screening panel to include DN107, DO100 and DV100, while retaining only those that performed well in the previous study as other four candidates. We applied no electric pulse as control for the last condition to build our screening panel consisting of 40 conditions with five buffers P1–P5 and eight electric pulse codes.

We activated polyclonal CD8^+^ T cells isolated from B6J mice *in vitro* using antibodies against CD3ε and CD28 for 48 hr before nucleofection. We then nucleofected activated CD8^+^ T cells with RNP complex containing three different CD90-targeting crRNAs (**Table 1**) and kept them in culture for additional two days in the presence of rmIL-2 but without anti-CD3ε/CD28 (**Figure 1A**). To determine the optimal condition, we evaluated the frequency of CD90^lo^ population (knockout efficacy) and cellular yield (calculated by dividing the recovered cell number by the number of cells seeded after nucleofection). Consistent with the notion that activated T cells are amenable to transfection and resultant CRISPR/Cas9-mediated genome editing (4), more than 90% of the harvested CD8^+^ T cells decreased the expression of CD90 protein under many of the nucleofection conditions (**Figure 1B, C, E**). In contrast, there was a sizable difference in the cellular yield among the tested nucleofection conditions, ranging from less than 10% to 150% or higher (**Figure 1D, E**).

**Figure 1.**
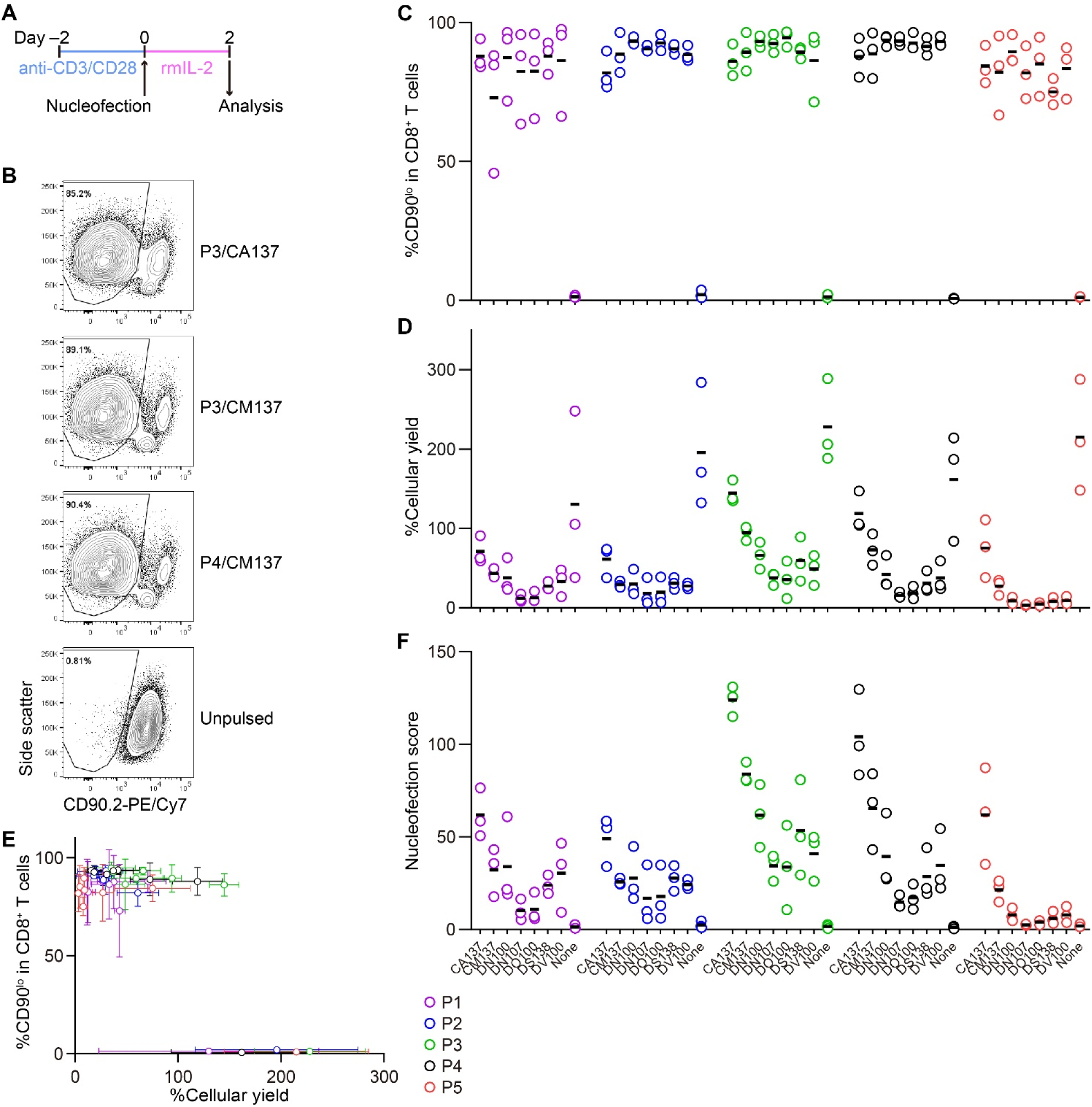
Assessment of knockout efficacy and cellular yield after CRISPR/Cas9 genetic engineering of *in vitro*-activated CD8^+^ T cells using nucleofection. (**A**) Experimental procedure. After activation with anti-CD3ε/CD28 antibodies for 2 days, cells were nucleofected with three CD90 crRNAs and then kept in culture in the presence of rmIL-2 for 2 days before analysis. (**B**) Representative plots from one experiment showing the downregulation of CD90 protein 2 days after nucleofection. Plots are gated on viable CD8^+^ T cells. (**C**–**D**) %CD90.2^lo^ in viable CD8^+^ T cells (**C**) and cellular yield (**D**) on day 2. (**E**) 2D-plot representation of the data shown in **C** and **D**. (**F**) Nucleofection score calculated as [%CD90^lo^] × [%Yield] / 100. Graphs show pooled data from three independent experiments.

To make an objective decision based on these two readouts, we derived a unified parameter, nucleofection score, from knockout efficacy and cellular yield, defined as ([%CD90^lo^] × [%Yield]) / 100. By definition, a condition that exhibits 100% knockout efficacy and 100% cellular yield gives a nucleofection score of 100, while the one with 0% knockout efficacy gives 0 irrespective of cellular yield. We found that two conditions, P3 buffer with the pulse code CA137 (P3/CA137) and P4/CA137, marked clearly higher mean score than others (**Figure 1F**). P3/CM137, P3/DN100, P4/CM137 and P5/CA137 yielded comparable nucleofection scores, which were slightly lower than P3/CA137 and P4/CA137. Based on these results, we decided to test the *in vivo* immune responsiveness of *in vitro*-activated CD8^+^ T cells nucleofected using P3/CA137, P3/CM137 and P4/CM137. We selected P4/CM137 from the above listed four conditions because this was what Seki and Rutz identified as the optimal condition (4).

### Effector and memory cell differentiation of *in vitro*-activated, nucleofected primary CD8^+^ T cells in HSV-OVA-infected host

To examine *in vivo* antigen-specific responses of nucleofected *in vitro*-activated CD8^+^ T cells, we nucleofected OT-I cells as we did for the screening using three CD90 crRNAs (**Table 1**). After letting the cells rest for 2 days in the presence of rmIL-2 (i.e. 4 days after activation), 150,000 each of the nucleofected cells were adoptively transferred into HSV-OVA-infected hosts (**Figure 2A**). Recipients were split into two groups, with each group receiving two populations of nucleofected OT-I cells (three with CD90 crRNAs and one no RNP control). To gauge the influence of nucleofection itself, we co-injected 150,000 OT-I cells into all recipients that were in the same manner as CD90-targeted cells but without nucleofection (denoted as NTC). Thus, each recipient received three populations of OT-I cells expressing GFP, tdT or CD45.1. Recipients were infected on the day when OT-I cells were put in culture for activation (i.e. 4 days before the adoptive transfer).

**Figure 2.**
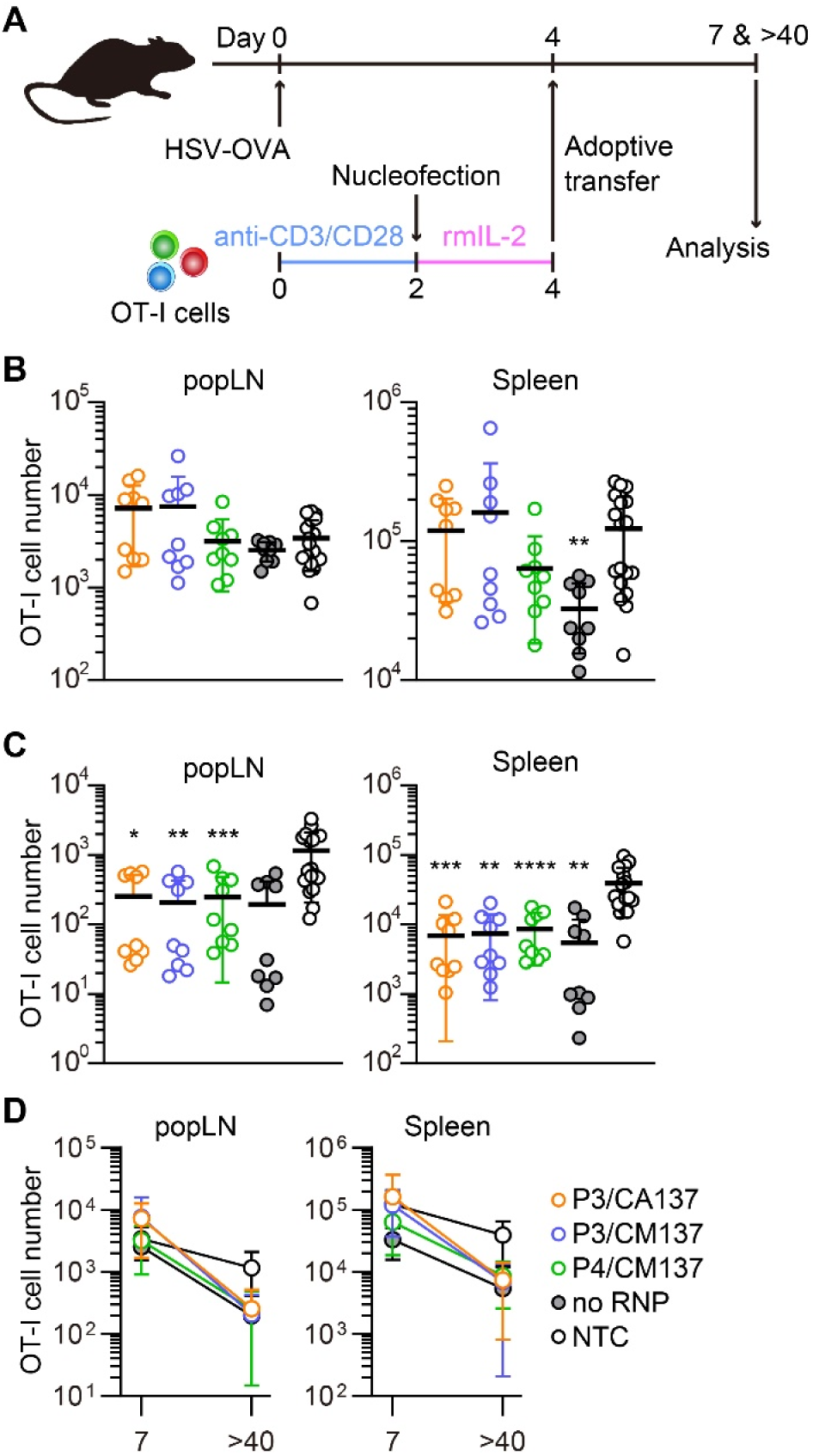
*In vivo* survival/expansion of nucleofected *in vitro*-activated CD8^+^ T cells. (**A**) Experimental scheme. OT-I cells were nucleofected with three CD90 crRNAs using P3/CA137, P3/CM137 or P4/CM137. After nucleofection and 2-day culture in the presence of rmIL-2, 150,000 OT-I cells per group were injected into hosts that were subcutaneously infected with HSV-OVA 4 days before adoptive transfer of OT-I cells. Each host received three groups including NTC control. (**B, C**) OT-I cell number in popLN and spleen on days 7 (**B**) and > 40 (**C**) after infection. (**D**) Data shown in **B** and **C** plotted in 2D to gauge the severity of contraction. Graphs show pooled data from two independent experiments with n = 8–9 or 14–15 each nucleofected groups or non-nucleofected control, respectively. Congenic marker assignment was swapped in each experiment. NTC, non-nucleofected control. \**p* < 0.05, \*\**p* < 0.01, \*\*\**p* < 0.001, \*\*\*\**p* < 0.0001 as compared to NTC by Kruskal-Wallis test with Dunn’s multiple comparison.

On days 7 and >40 post-infection, we determined the number of OT-I cells in antigen-draining popLN and spleen. Although there was a weak trend on day 7 that OT-I cells nucleofected using P3 buffer outperformed other groups (**Figure 2B**), none of the nucleofected OT-I cell populations displayed a statistically significant difference from NTC OT-I cells, except that no RNP control in the spleen was slightly less abundant. After >40 days, the number of nucleofected OT-I cells were less abundant for all groups in both popLN and spleen as compared to NTC OT-I cells, resulting in a subtly more pronounced contraction of nucleofected OT-I cells (**Figure 2C, D**). Longitudinal analysis of OT-I cells among PBL revealed that the severer contraction of nucleofected cells manifested after day 21 post-infection (**Supplementary Figure 1A**). Frequency of CD90^lo^ cells among OT-I cells remained high in all three test groups, while no RNP control OT-I cells retained CD90 expression at the same frequency as NTC OT-I cells (**Supplementary Figure 1B–E**). Thus, the loss of CD90 did not account for the greater degree of contraction of nucleofected OT-I cells.

To further characterize *in vitro*-activated, nucleofected OT-I cells, we analyzed their phenotype after adoptive transfer into HSV-OVA-infected host. On day 7 post-infection, most OT-I cells exhibited CD127^+^ KLRG1^−^ memory precursor effector cell phenotype (**Figure 3A, Supplementary Figure 2A**), with CD127^−^ KLRG1^−^ early effector cells being the second most abundant population. Such composition of effector OT-I cells in popLN and spleen was common to all five conditions (and hence irrespective of the loss of CD90). Yet, nucleofected OT-I cells contained higher frequency of memory precursor effector cells than NTC OT-I cells did.

**Figure 3.**
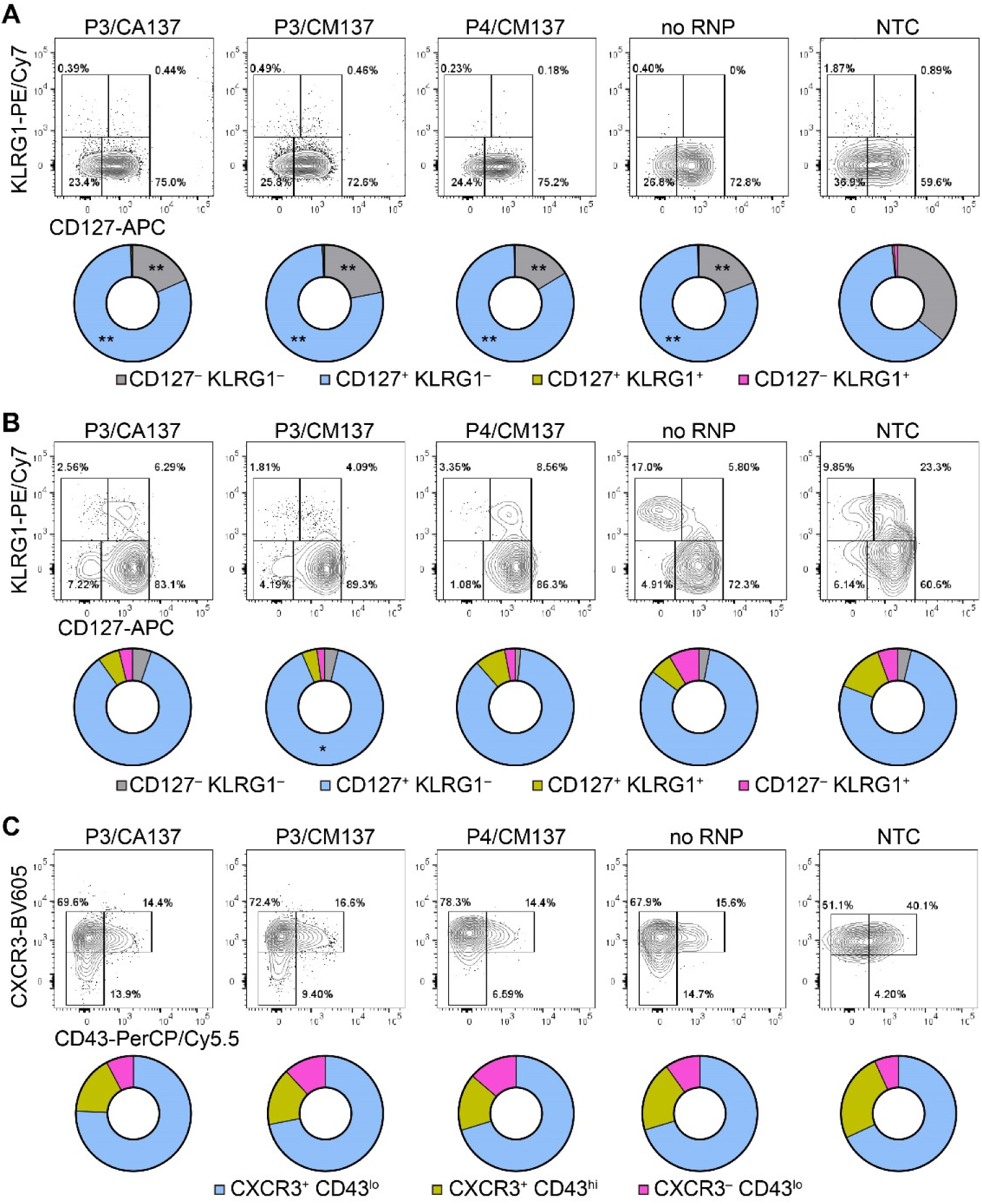
Phenotype of *in vitro*-activated OT-I cells in spleen after adoptive transfer into HSV-OVA-infected hosts. OT-I cells were nucleofected as in **Figure 2**. (**A, B**) Expression of CD127 and KLRG1 on OT-I cells in the spleen on days 7 (**A**) and > 40 (**B**). Pie charts show the mean frequencies of four populations identified by these two markers. (**C**) Expression of activation-associated glycoform of CD43 and CXCR3 on OT-I cells in the spleen > 40 days after infection. Pie charts show the mean frequencies of three populations identified by these two markers. Graphs show pooled data from two independent experiments with n = 9 or 15 for each nucleofected group or non-nucleofected control, respectively. Flow cytometric plots are gated on viable OT-I cells identified by the expression of congenic markers and show concatenated data from one of two experiments with n = 5 per group. Congenic marker assignment was swapped in each experiment. NTC, non-nucleofected control. \**p* < 0.005, \*\**p* < 0.01 as compared to NTC by ordinary two-way ANOVA with Dunnett’s multiple comparison.

CD127^+^ KLRG1^−^ cells remained the dominant population until > 40 days post-infection, with small fractions of cells being CD127^−^ KLRG1^−^ CD127^+^ KLRG1^+^ or CD127^−^ KLRG1^+^ (**Figure 3B, Supplementary Figure 2B**). As a result, all five groups generated memory cells with a largely comparable composition as assessed by the expression of CD127 and KLRG1. To confirm that memory OT-I cell heterogeneity is indeed comparable, we also evaluated the expression of activation-associated glycoform of CD43 (recognized by the antibody clone 1B11) and CXCR3. The majority of OT-I cells in all five groups comprised CXCR3^+^ CD43^lo^ cells (**Figure 3C, Supplementary Figure 2C**), which had been reported to self-renew at an intermediate rate but exhibit greater clonal expansion upon recall responses (19). There was a weak trend that NTC OT-I cells contained slightly higher frequency of CXCR3^+^ CD43^hi^ cells, but the difference did not reach a statistically significant level. Even though there were small differences in the phenotype between nucleofected and NTC OT-I cells, none of them can explain the lower number of memory cells generated by nucleofected OT-I cells (**Figure 2C, D**). Furthermore, nucleofected OT-I cells displayed subtle differences in cytokine producing ability on day 7 as compared to NTC OT-I cells (**Supplementary Figure 3**), whereas there was a slight decrease in the frequency of IFN-γ^+^ cells (but not IFN-γ^+^ TNF-α^+^ cells) >40 days after infection (**Supplementary Figure 4**).

In sum, nucleofection could affect the fitness of *in vitro*-activated CD8^+^ T cells, without reshaping the differentiation spectrum of effector and memory CD8^+^ T cells. Based on these results, we concluded P3/CA137 as the best condition for CRISPR/Cas9-mediated gene inactivation of *in vitro*-activated primary mouse CD8^+^ T cells because it marked the highest nucleofection score among the tested conditions (**Figure 1F**). In addition, the observation that inactivation of CD90 did not lead to any functional alteration as compared to no RNP control confirmed its suitability for use in our screening and functional validation.

### CRISPR/Cas9-mediated genetic engineering of naïve primary mouse CD8^+^ T cells

Next, we tested the optimal nucleofection conditions for naïve primary mouse CD8^+^ T cells. Similar to the screening for *in vitro*-activated CD8^+^ T cells, we selected seven electric pulse codes based on the previous publication (4) and inputs from the manufacturer: CM137, DN100, DP100, DS137, DS150, DZ100 and EA100. Of these, DP100, DZ100 and EA100 were not included in the previous study (4). With the no pulse control for each buffer, we evaluated again 40 conditions.

CD8^+^ T cells were nucleofected with three CD90 crRNAs (**Table 1**) after incubating with rmIL-7 for 24 hr and analyzed for the loss of CD90 protein and cellular yield on day 5 post-nucleofection (**Figure 4A**). Unlike *in vitro*-activated CD8^+^ T cells, naïve CD8^+^ T cells lost the expression of CD90 protein to a variable degree, ranging from less than 60% to approximately 90% on average (**Figure 4B, C, E**). On top of that, cellular yield was much lower than that of activated CD8^+^ T cells (**Figure 4D, E**). To objectively rank all the tested conditions, we used the nucleofection scores again and found that P4/CM137 and P5/CM137 exhibited slightly higher score than other conditions (**Figure 4F**), owing largely to higher cellular yield. The previously reported optimal condition, P4/DS137, was one of the conditions with the highest knockout efficacy but ranked only fifth in our screening because of the lower cellular yield as compared to P4/CM137 and P5/CM137. P1/CM137 and P2/CM137, the third and fourth position in our result, yielded marginally higher score than P4/DS137 because of higher cellular yield. Yet these two conditions achieved less than 80% knockout efficacy, which is lower than that of P4/DS137 at 88.9%. Considering that higher knockout efficacy is more beneficial than slightly higher cellular yield for most experiments, we chose P4/CM137, P4/DS137 and P5/CM137 to evaluate the immune responsiveness of nucleofected naïve CD8^+^ T cells.

**Figure 4.**
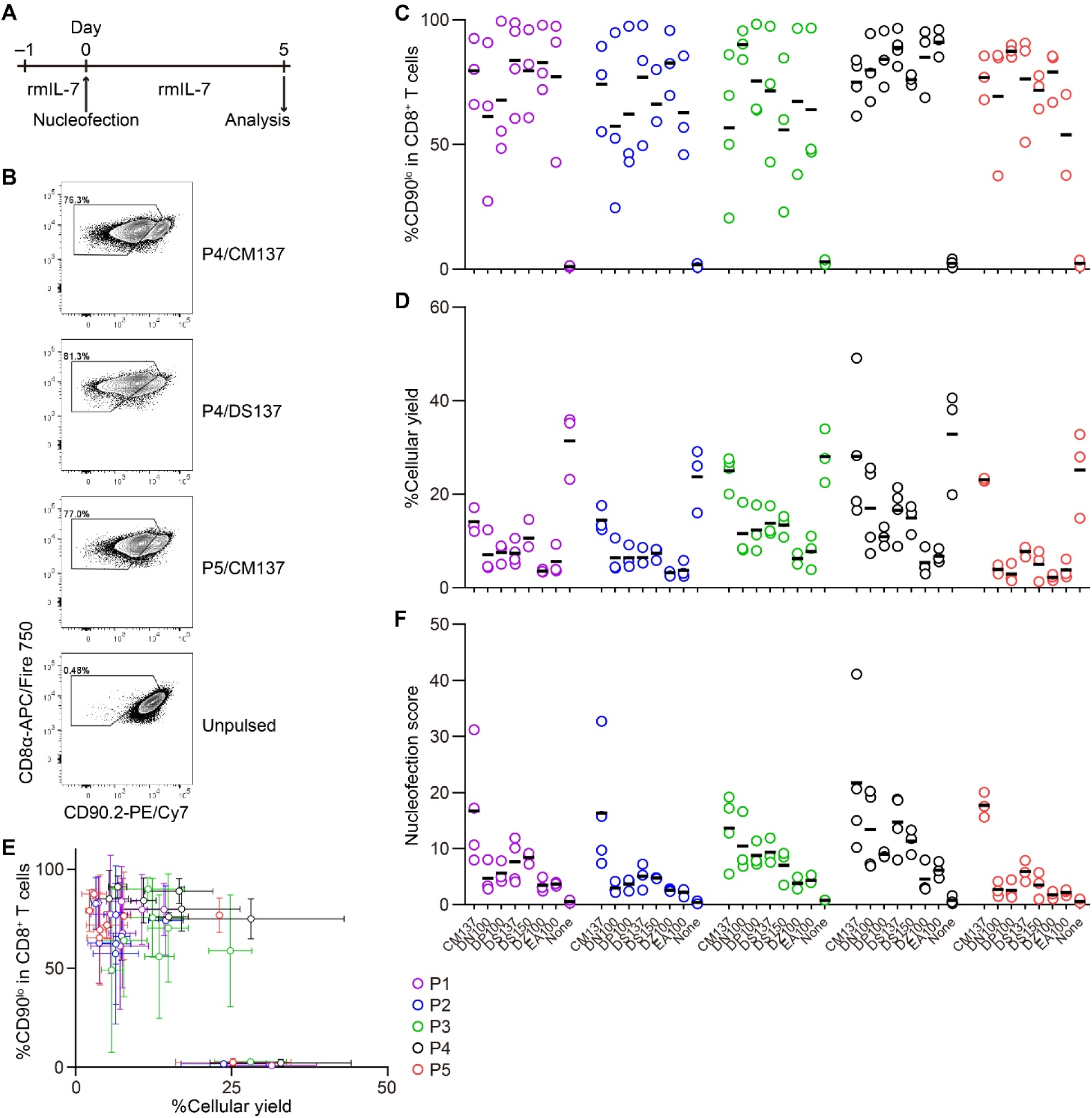
Assessment of knockout efficacy and cellular yield after CRISPR/Cas9 genetic engineering of naïve CD8^+^ T cells using nucleofection. (**A**) Experimental procedure. Magnetically isolated CD8^+^ T cells were kept in culture in the presence of rmIL-7 for 24 hr before nucleofection. Nucleofected cells were kept in culture for additional 5 days before analysis. (**B**) Representative plots from one experiment showing the downregulation of CD90 protein 5 days after nucleofection. Plots are gated on viable CD8^+^ T cells. (**C, D**) %CD90.2^lo^ in viable CD8^+^ T cells (**C**) and cellular yield (**D**) on day 2. (**E**) 2D-plot representation of the data shown in **C** and **D**. (**F**) Nucleofection score calculated as [%CD90^lo^] × [%Yield] / 100. Graphs show pooled data from 3–4 independent experiments.

### Antiviral responses of nucleofected primary naïve CD8^+^ T cells

To test *in vivo* immune responses of nucleofected naïve CD8^+^ T cells, we nucleofected OT-I cells with three CD90-targeting crRNAs using the above selected three conditions and cultured for 5 days in the presence of rmIL-7 (**Figure 5A**). One day before subcutaneous infection with HSV-OVA, we injected 5,000 each of nucleofected OT-I cells into recipient mice. Similar to the assay for in vitro-activated CD8^+^ T cells, we split recipients into two groups and injected two groups of nucleofected OT-I cells (two of P4/CM137, P4/DS137, P5/CM137 or no RNP control) per recipient, together with freshly-isolated OT-I cells as a third group as benchmark. Thus, each recipient received a total of 15,000 OT-I cells, which were identifiable by the expression of GFP, tdT or CD45.1.

**Figure 5.**
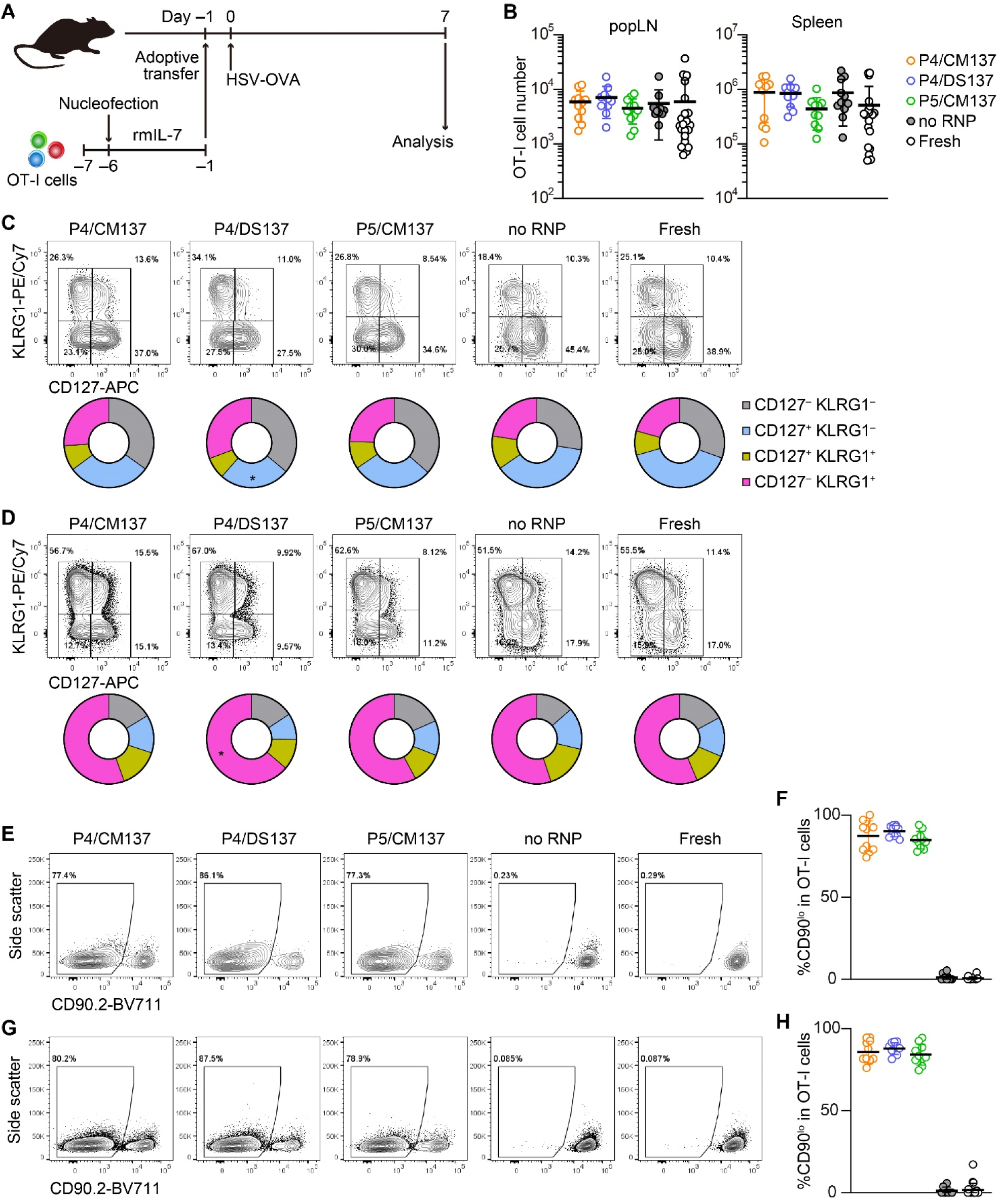
*In vivo* antiviral immune response of nucleofected naïve CD8^+^ T cells. (**A**) Experimental scheme. OT-I cells were nucleofected with three CD90 crRNAs using P4/CM137, P4/DS137 or P5/CM137. One day before subcutaneous HSV-OVA infection, 5,000 OT-I cells per group were injected into hosts. Each recipient received three groups including freshly isolated OT-I cells. (**B**) OT-I cell number on day 7 post-infection. No statistically significant difference between nucleofected groups and freshly isolated OT-I cells by Kruskal-Wallis test with Dunn’s multiple comparison (popLN) or ordinary one-way ANOVA with Dunnett’s multiple comparison (spleen). (**C, D**) CD127 and KLRG1 expression on OT-I cells in popLN (**C**) and spleen (**D**) on day 7. \**p* < 0.05 by ordinary two-way ANOVA with Dunnett’s multiple comparison. (**E–H**) Frequency of CD90^lo^ cells in OT-I cells in popLN (**E, F**) and spleen (**G, H**) on day 7. Flow cytometric plots are gated on viable OT-I cells identified by the expression of congenic markers and show concatenated data from one of three experiments with n = 5 per group. Congenic marker assignment was swapped in each experiment. Graphs show pooled data from three independent experiments with n = 10–11 or 21–22 for nucleofected groups and freshly-isolated cells, respectively.

The number of nucleofected OT-I cells day 7 post-infection was indistinguishable from that of freshly-isolated OT-I cells in both popLN and spleen (**Figure 5B**). In all five groups, OT-I cells in popLNs exhibited a diverse effector cell differentiation profile identified by the expression of CD127 and KLRG1 (**Figure 5C**). In contrast, the majority of OT-I cells in the spleen displayed CD127^−^ KLRG1^+^ short-lived effector cell phenotype, irrespective of the condition (**Figure 5D**). After undergoing massive clonal expansion, OT-I cells nucleofected with CD90-targeting RNPs maintained the frequency of CD90^lo^ cells similar to the level found in the screening, whereas mock-nucleofected and freshly-isolated OT-I cells retained the high expression of CD90 (**Figure 5E–H, Figure 4C**). These results indicate that loss of CD90 before priming affected neither clonal expansion nor heterogeneity of effector OT-I cells. Taken together, once CD8^+^ T cells survive nucleofection, they are as competent as freshly-isolated CD8^+^ T cells to elicit effector responses. Although P4/CM137 and P5/CM137 marked slightly higher nucleofection scores (**Figure 4F**), we concluded P4/DS137 as the best condition for naïve primary mouse CD8^+^ T cells based on its higher knockout efficacy (**Figure 4C**).

### Co-targeting of second gene for enrichment of successfully gene-edited cells

Our optimal nucleofection conditions constantly yielded > 75% knockout efficacy even for naïve CD8^+^ T cells (**Figure 4B, C** and **Figure 5E-H**) that were more resistant to gene inactivation than *in vitro*-activated cells. However, use of such “partial knockout” cells conceivably imposes a hurdle for interpretation of experimental results when the experiment aims to identify functions of the gene of interest. For example, residual wild-type cells can outnumber knockout cells or affect their behavior *in vivo*. Thus, it is highly useful to devise a tool to enrich successfully targeted cells to a purity of 90% or higher even when the knockout efficacy does not reach such a high level after nucleofection. For targets that are expressed uniformly on the surface of naïve CD8^+^ T cells (e.g. CD90), isolation of successfully targeted cells is readily attainable by simple antibody staining followed by cell sorting. Yet this is not the case for intracellular proteins (e.g. transcription factors) or surface proteins that are completely or partially absent on normal naïve CD8^+^ T cells (e.g. PD-1). Therefore, we tested whether a simultaneous targeting of a “reporter” gene enables to identify and enrich successfully targeted cells. To this end, we chose the intracellular protein DOCK2 as the model target because a reporter mouse line that expresses a GFP-fused form of DOCK2 (16) allows for single-cell analysis of its expression levels by flow cytometry. We used CD90 as the reporter because of its uniform expression on naïve CD8^+^ T cells and the lack of functional impact on them (**Figure 5**). To assess the effectiveness of enrichment across a wide range of knockout efficacy, we deliberately varied knockout efficacy of DOCK2 by using different combinations of crRNAs. We kept the knockout efficacy for CD90 low by decreasing the amount of RNP complex to deliver into CD8^+^ T cells, such that the frequency of CD90^lo^ cells always remains lower than that of DOCK2-GFP^lo^ cells.

We followed changes in the frequency of DOCK2-GFP^lo^ cells among total, CD90^hi^ and CD90^lo^ CD8^+^ T cells up to 10 days post-nucleofection. In most cases, the frequency of DOCK2-GFP^lo^ and CD90^lo^ among total CD8^+^ T cells rapidly increased by day 5 and remained relatively stable thereafter. CD90^lo^ cells were indeed enriched for DOCK2-GFP^lo^ cells as compared to total CD8^+^ T cells at all timepoints in all experiments (**Figure 6A–F**). We confirmed lower expression levels of *Dock2* mRNA among CD90^lo^ cells as compared to CD90^hi^ cells (**Figure 6G**). To evaluate the effectiveness of simultaneous CD90 targeting as the reporter, we calculated %enrichment as ([%DOCK2-GFP^lo^ in CD90^lo^ cells] – [%DOCK2-GFP^lo^ in total CD8^+^ T cells]) / (100 – [%DOCK2-GFP^lo^ in total CD8^+^ T cells]). This value shows to what extent the use of reporter filled the gap for the observed %DOCK2-GFP^lo^ among total CD8^+^ T cells to reach 100% knockout efficacy, without being affected by deliberately varied %DOCK2-GFP^lo^. The mean %enrichment values on days 5, 7 and 10 were almost identical at approximately 50 (**Figure 6H**), indicating that enrichment efficacy does not depend on the time after nucleofection. Furthermore, these data demonstrate that the enrichment using CD90 as reporter does not always yield > 90% purity of DOCK2^lo^ cells. Of note, within the tested ranges of knockout efficacy for DOCK2 and CD90, %enrichment positively correlated with %CD90^lo^ and %DOCK2-GFP^lo^ cells among total CD8^+^ T cells (**Figure 6I, J**), where %DOCK-GFP^lo^ showed better correlation than %CD90^lo^ did. Taken together, simultaneous targeting of a reporter gene can provide a means to enrich the cells that underwent successful editing of the primary target gene, albeit to a limited degree.

**Figure 6.**
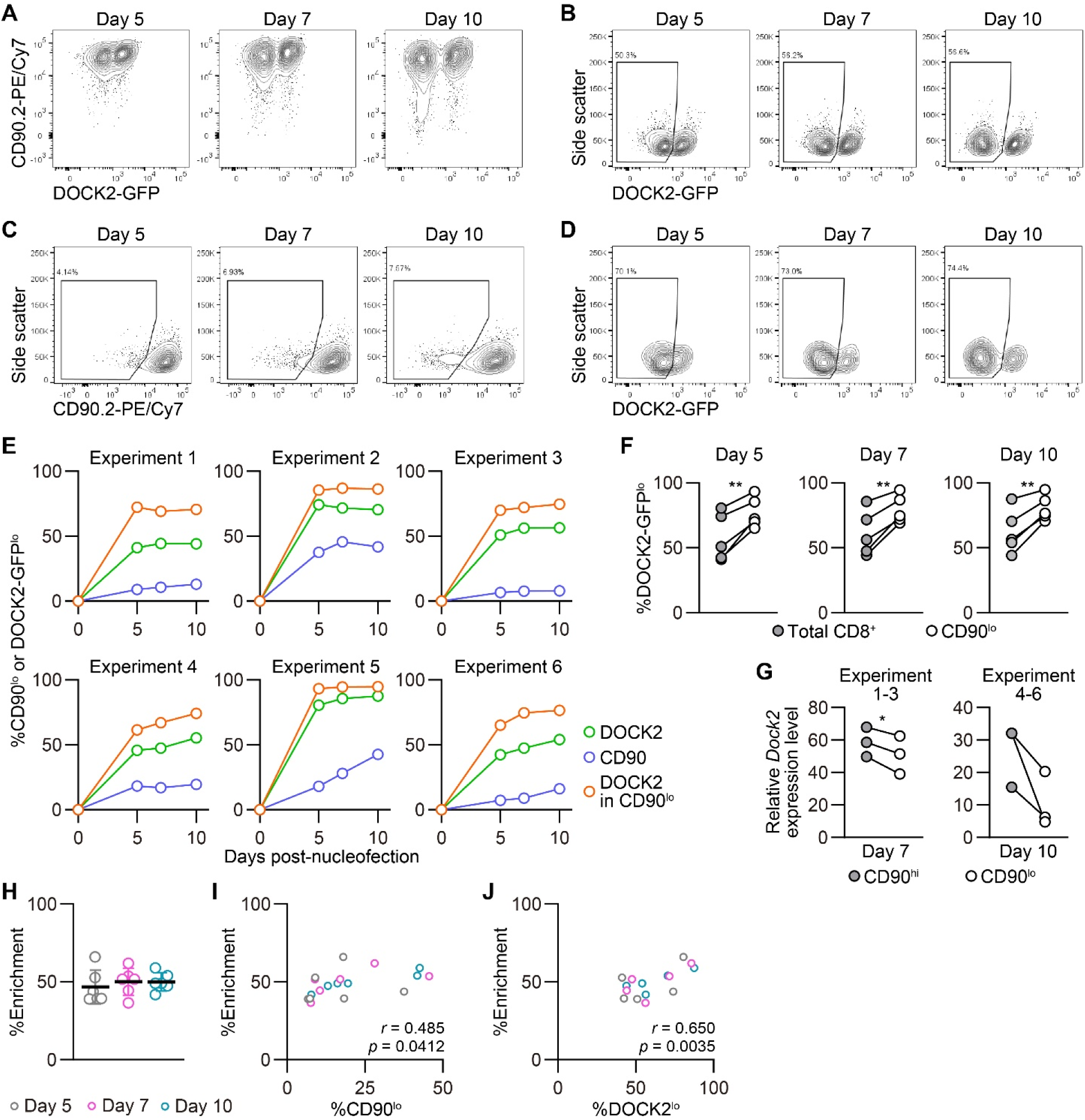
Enrichment of DOCK2-knockout cells using simultaneously targeted CD90 as a reporter. DOCK2-GFP CD8^+^ T cells were nucleofected with CD90 and/or DOCK2 crRNAs using P4/DS137. (**A–D**) Representative flow cytometric plots of DOCK2-GFP versus CD90 (**A**), DOCK2-GFP (**B**) and CD90 (**C**) among total CD8^+^ T cells and DOCK2-GFP among CD90^lo^ cells (**D**). Plots are taken from experiment 3. (**E**) %CD90^lo^ cells among total CD8^+^ T cells and %DOCK2-GFP^lo^ cells among total and CD90^lo^ CD8^+^ T cells. (**F**) Paired comparison of %DOCK2-GFP^lo^ cells between total and CD90^lo^ CD8^+^ T cells in six independent experiments. \*\**p* < 0.005 by paired *t*-test. (**G**) *Dock2* mRNA level in sorted CD90^lo^ and CD90^hi^ CD8^+^ T cells. Data are normalized to the expression level of *Dock2* in the control cells that are nucleofected only with three CD90 crRNAs. Cells were sorted on day 7 or 10 for experiments 1–3 or 4–6, respectively. \**p* < 0.05 by paired *t*-test. (**H**) Enrichment efficacy of DOCK2-GFP^lo^ cells by gating on CD90^lo^ cells on each time point measured with %enrichment values. No statistically significant difference by Kruskal-Wallis test with Dunn’s post-hoc. (**I, J**) Correlation between %enrichment and the frequency of CD90^lo^ (**I**) or DOCK2-GFP^lo^ cells (**J**) among total viable CD8^+^ T cells. Pearson correlation coefficient was computed using the pooled data of six independent experiments.

### Disparate decay rate of DOCK2 and CD90 after genome editing

The relatively stable frequency of CD90^lo^ and DOCK2^lo^ cells (**Figure 6E**) prompted us to ask whether the loss of DOCK2 and CD90 protein after gene inactivation would also progress at a comparable rate. To answer this question, we traced changes in the mean fluorescence intensity (MFI) of DOCK2-GFP and CD90 among successfully targeted cells by flow cytometry. To take into account inter-experimental variabilities in the staining, we obtained normalizers for each experiment by dividing the measured MFI value of non-targeted control by the mean values of non-targeted control from six experiments. Then, all the measured MFI values were normalized using the normalizer of corresponding experiment. Interestingly, despite the deliberate variability in the frequency of knockout cells, normalized MFI values turned out to decay at a comparable rate in all the experiments, as shown by the extremely small variability (**Figure 7A**). We found that the MFI values of both DOCK2-GFP among DOCK2-GFP^lo^ cells and CD90 among CD90^lo^ cells kept decreasing until day 10 (**Figure 7A**). To estimate the half-life of these proteins after nucleofection, we calculated relative MFI by normalizing the measured values in each experiment to that of the corresponding non-targeted cells. By fitting the data to a one phase decay model, we obtained the half-life of 2.57 and 1.53 days (95% CI 1.88–3.51 and 1.30–1.77) for DOCK2 and CD90, respectively (**Figure 7B**), showing that the decay rate of DOCK2 is slower than that of CD90. In contrast, changes in %DOCK2-GFP^lo^ and %CD90 from day 5 onwards appeared comparable and smaller than those between day 0 and 5 (**Figure 6E**). Thus, we hypothesized that gene inactivation in successfully nucleofected cells was largely completed within 5 days after nucleofection for both targets. To test this, we assumed the frequency of knockout cells on day 10 as plateau of each experiment and normalized the frequency of knockout cells on days 5 and 7 to that on day 10 (**Figure 7C**). After curve fitting, we obtained the half-time to reach the putative plateau of 1.48 and 1.80 days (95% CI 1.74–2.50 and 1.97 to 3.17) for DOCK2 and CD90, respectively. These results indicate that the loss of protein proceeds at a disparate rate for different proteins, due presumably to variable turnover rate and abundance of each protein. Yet the inactivation of genes become completed within approximately the same time. Moreover, extremely small variability in the normalized MFI values (**Figure 7A**) implies that degradation of protein (and mRNA) encoded by the target gene occurs at a rate that is dependent on its biochemical properties of the target but independent of the knockout efficacy.

**Figure 7.**
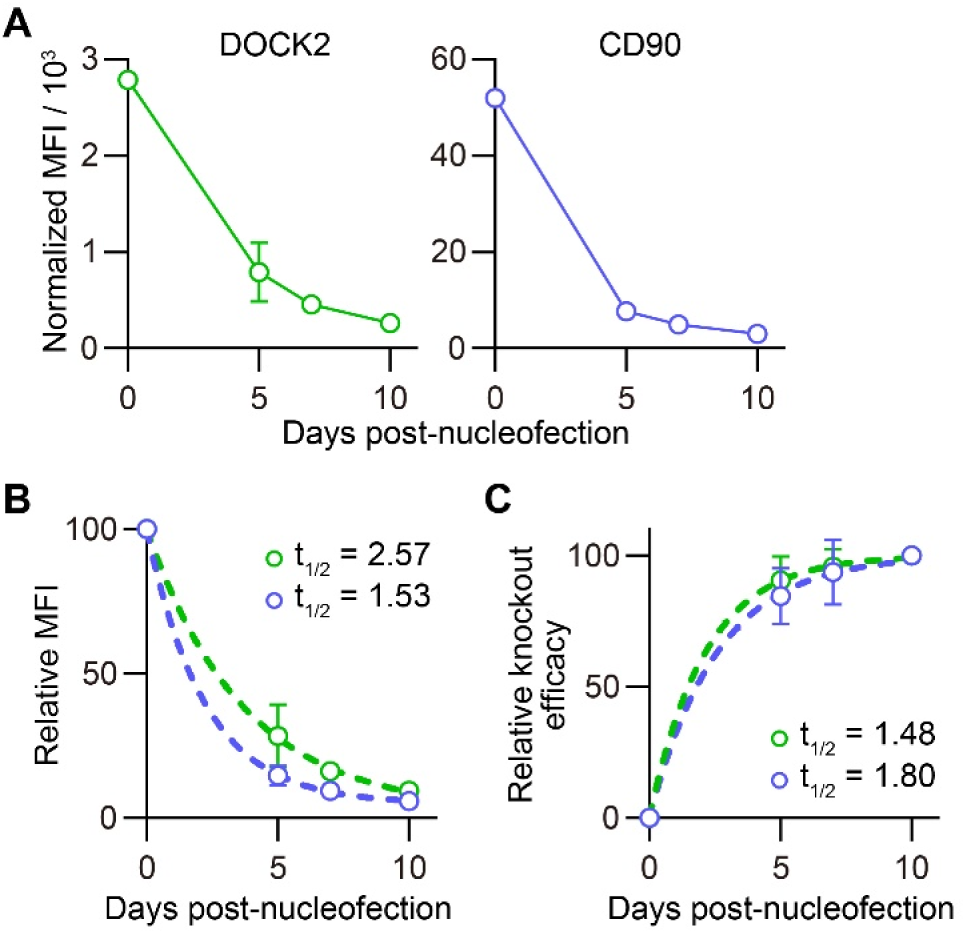
Disparate kinetics of gene inactivation and protein loss for DOCK2 and CD90. (**A**) Decay of normalized MFI values of DOCK2-GFP and CD90 among DOCK2-GFP^lo^ and CD90^lo^ cells, respectively. (**B**) Relative MFI values of DOCK2-GFP and CD90 proteins calculated by normalizing the values presented in (**A**) to the MFI value on day 0. Curves defined as y = (100 – [Plateau]) × exp(– K × x) were fitted to the data, with *R*^*2*^ = 0.979 and 0.997 for DOCK2 and CD90, respectively. Plateau values and half-life were computed during curve fitting. (**C**) Kinetics of relative knockout efficacy calculated by normalizing %DOCK2-GFP^lo^ and %CD90^lo^ in each experiment to the corresponding values on day 10. Data were fitted with the equation y = [Plateau] × (1 – exp(–K×x)), with *R*^*2*^ = 0.984 and 0.970 for DOCK2 and CD90, respectively. Plateau value and half-time to reach the plateau was estimated from the data during curve fitting. Graphs show pooled data from six independent experiments, except that values of %CD90^lo^ from experiments 5 and 6 (as indicated in **Figure 6**) were excluded in **C** because of their continuous increase until day 10.

## Discussion

In this study, we determined optimal conditions for CRISPR/Cas9-mediated genetic engineering of primary mouse CD8^+^ T cells using nucleofection. We found that naïve CD8^+^ T cells retained their ability to elicit *in vivo* antiviral responses after nucleofection and 5-day long culture in the presence of rmIL-7. Although *in vitro*-activated CD8^+^ T cells displayed slightly impaired *in vivo* expansion and/or survival in virus-infected hosts, they displayed a comparable status of effector and memory CD8^+^ T cells. Thus, nucleofection-based CRISPR/Cas9 genome editing is a fast and efficient approach to generate mutant cells from primary CD8^+^ T cells without degenerating their functions.

Congenic CD90 has been widely used for adoptive transfer experiments (20–22) and as a non-fluorescent reporter of genes that encode intracellular proteins or those that are difficult to label using antibodies (23–27). Yet the function of CD90 in CD8^+^ T cells remains elusive, with contradictory findings. Antibody blockade of CD90 during *in vitro* T cell activation appears to promote proliferation and acquisition of the expression of effector molecules (28). Similarly, CD90-deficient T cells display impaired phosphorylation of the tyrosine kinase Lck and attenuated calcium mobilization following anti-CD3ε stimulation, resulting in attenuated proliferation and delayed hypersensitivity reaction (29). On the contrary, another study reported that CD90 deficiency augments TCR signaling in thymocytes, which in turn leads to lower single-positive thymocyte output from double-positive thymocytes (30). Our results provided yet another view that loss of CD90 in mature T cells induced by CRISPR/Cas9 genetic engineering does not alter the *in vivo* immune responsiveness of naïve CD8^+^ T cells. This is consistent with the previous finding that the proliferative response of CD90^lo^ and CD90^hi^ hemagglutinin-specific TCR-transgenic CD8^+^ T cells isolated from 18–20 months-old mice is indistinguishable after *in vitro* stimulation with their cognate peptide (31). A potential explanation for such a discrepancy is that the absence of CD90 during T cell development may imprint a minor functional defect, while it has less significant roles in mature T cells. Furthermore, the fact that nucleofected CD90-proficient and deficient CD8^+^ T cells responded similarly to antigenic stimulation demonstrates that neither the nucleofection process nor transient presence of gRNA-Cas9 RNP affects CD8^+^ T cell functions. In contrast to nucleofection-based delivery of CRISPR/Cas9 machinery, viral transduction forces long-lasting expression of gRNA, Cas9 and other virus-derived gene products, which can increase the risk of rejection after adoptive transfer of engineered T cells. Since RNP complex disappears over time after nucleofection, there is conceivably much less chance of the rejection with nucleofection as compared to viral transduction.

We observed that the simultaneous targeting of a second gene as reporter allows for partial enrichment of cells that are successfully engineered at primary target loci. One of the reasons that motivated us to test this was the fact that several clinical studies suffered from poor knockout efficacy. For example, attempts to generate *CCR5*-deficient bone marrow cells (32) and *PDCD1*-deficient T cells (33) proved the safety of cell-based therapies using CRISPR/Cas9-engineered cells. However, the knockout efficacy in these studies remained 50% at the highest (32–34). Albeit less frequent, such a scenario where the knockout efficacy does not reach the desirable level can also occur in basic research. Our enrichment strategy may help improve the knockout efficacy when the knockout efficacy already lies above 70–80%. Yet, we postulate that knock-in-knockout approaches by introducing a reporter gene into the target locus through homology-directed repair (34, 35) will serve a more stringent method to isolate successfully targeted cells, especially when it is combined with optimal crRNA design and Cas nuclease.

Proteomic analysis by other groups showed that CD90 is one of the most abundant proteins in mouse CD8^+^ T cells, with its copy number being about 9-fold higher than that of DOCK2 (36, 37). Yet we found that depletion of DOCK2 protein after gene inactivation takes longer time than that of CD90. The initial abundance of target proteins is undoubtedly an important factor that determines the duration required for its depletion after gene inactivation. Our results suggest nevertheless that the decay of target protein greatly depends on its physiological turnover rate as well. In fact, a recent study on the turnover rate of proteins in human CD8^+^ T cells revealed a large variability of protein half-life, ranging from less than a day to more than ten days (38). For example, less than 10% of human DOCK2 protein is replaced within 24 hr, making DOCK2 one of the most slowly renewing proteins. Although precise turnover rate of DOCK2 and CD90 in normal mouse CD8^+^ T cells remains unknown, our observation that DOCK2 has longer half-life than CD90 after nucleofection seems consistent with the previous observation. Thus, it is important to adapt the post-nucleofection incubation period for each target protein to obtain a desirable level of its depletion. Optimal experimental design has to take both initial abundance and turnover rate of the target protein into consideration because these parameters are different for each protein and change depending on the activation and differentiation status of CD8^+^ T cells (36, 38). This is particularly relevant to study functions of the gene of interest in naïve CD8^+^ T cells or early after their activation, where cell division does not facilitate the depletion of targeted protein. It is worth noting that post-nucleofection incubation did not affect the *in vivo* immune response of genetically engineered CD8^+^ T cells, justifying to allow for such an interval to achieve desirable gene inactivation and depletion of its product.

A recent study showed that nucleofection rapidly activates p53 pathway in mouse memory CD8^+^ T cells, impairing their recall response *in vivo* after adoptive transfer (9). This is in stark contrast to what we observed for naïve CD8^+^ T cells, even though both are resting cells. Consistent with our results, another recent study showed that nucleofected naïve CD8^+^ T cells retain *in vivo* responsiveness against viral infection (10). One of the possible explanations for such a striking difference may be that naïve and memory CD8^+^ T cells have distinct sensitivity of p53 pathway activation upon DNA damages. Importantly, authors of these two studies injected nucleofected cells without culture, followed by viral infection immediately (10) or three days later (9). Thus, the interval between nucleofection and immune stimulation is unlikely to be a decisive factor causing the abortive response of nucleofected memory CD8^+^ T cells. Albeit much less likely, another potential cause of the different behavior of naïve and memory CD8^+^ T cells may lie in the use of different equipment in the studies on naïve and memory CD8^+^ T cells (9, 10). Understanding of the exact mechanism that impairs the response of nucleofected memory CD8^+^ T cells is relevant for the use of electroporation-based CRISPR/Cas9-mediated genetic engineering both in basic research and clinical use.

Direct engineering of primary CD8^+^ T cells enables to generate mutant cells lacking one or more genes within two weeks. This requires far shorter time than establishing a mutant mouse line, which can easily take months. In addition, generation of mutant T cells without interbreeding multiple mutant and congenic mice reduces the number of animals used to obtain compound mutant mouse lines, cohering well with the 3R principle. Although genetically modified animals and viral transduction remain as a vital tool in immunology, CRISPR/Cas9-based genetic engineering of primary mouse CD8^+^ T cell will serve as a versatile and less time-consuming alternative or add-on. To take full advantage of CRISPR/Cas9-technology, the biochemical properties of target gene product are critical factors to consider, besides the choice of crRNAs and Cas9. While our study focused on CD8^+^ T cells, we envision that nucleofection-based CRISPR/Cas9 genetic engineering of other immune cells will require similar considerations to achieve maximal efficacy.

## Author Contributions

PP and JA performed and analyzed experiments with help from LY. JA supervised research and wrote the manuscript with input from all co-authors.

## Acknowledgments

We thank Prof. David Hoogewijs and Prof. Barbara Rothen for allowing us to use their equipment, Cell Analytics Facility of the University of Fribourg for operational supports, Prof. Jens V Stein for continued support, and Danièla Grand and Antoinette Hayoz for excellent technical assistance.

## Supplementary Figures

**Supplementary Figure 1.**
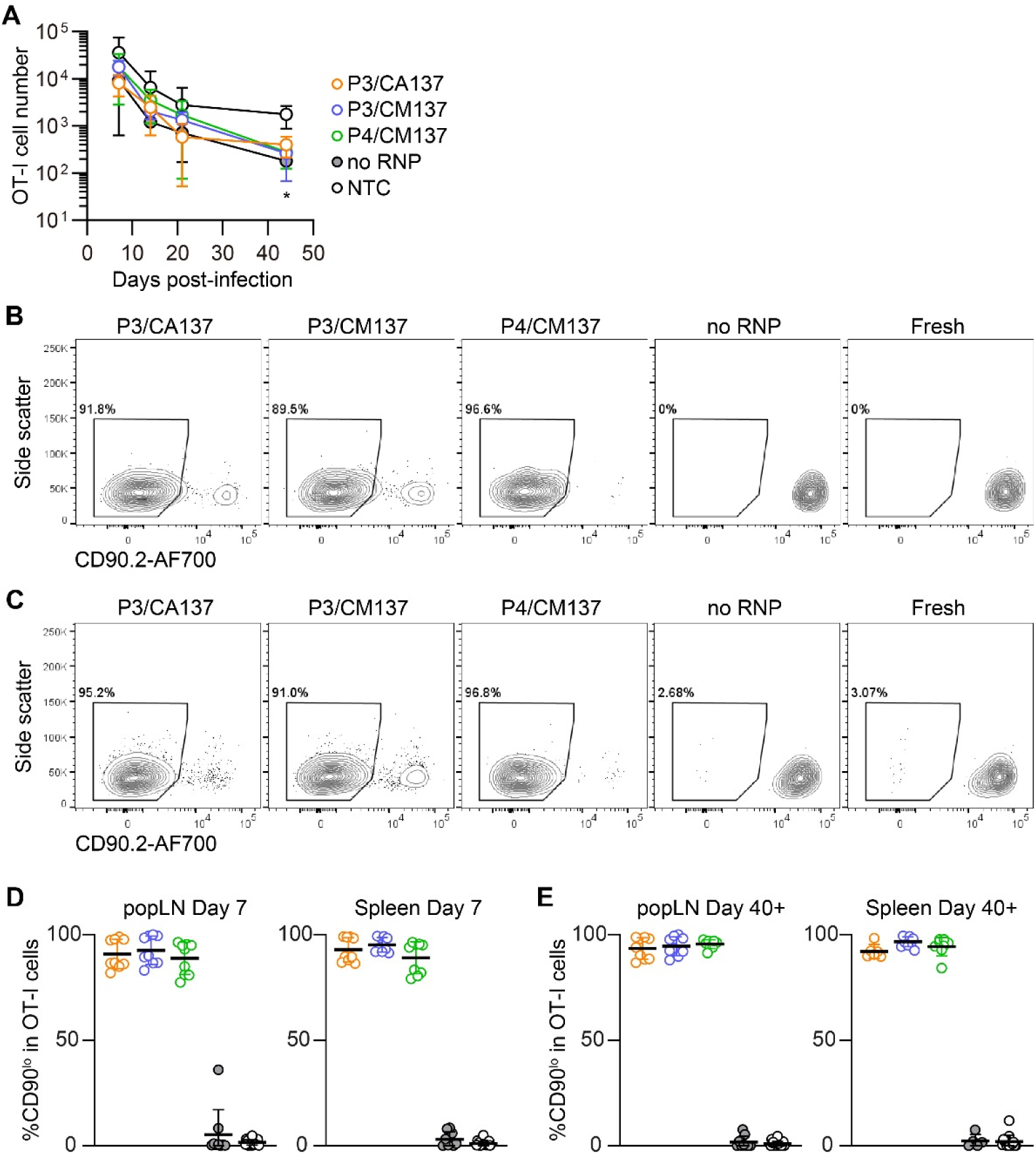
Kinetics of the number and CD90 expression of *in vitro*-activated OT-I cells after adoptive transfer into HSV-OVA-infected hosts. Nucleofection of OT-I cells and viral infection were performed as in **Figure 2**. (**A**) Kinetics of OT-I cell number per 1 mL blood. Statistical significance of differences between each of nucleofected group and control was analyzed by ordinary two-way ANOVA with Dunnett’s multiple comparison. \**p* < 0.0001. (**B–E**) Frequency of CD90^lo^ cells among OT-I cells in popLN and spleen on days 7 (**B, D**) and > 40 (**C, E**). Congenic marker assignment was swapped in each experiment. Data are pooled from two independent experiments with n = 6–9 or n = 12–15 for nucleofected cells or non-nucleofected control, respectively.

**Supplementary Figure 2.**
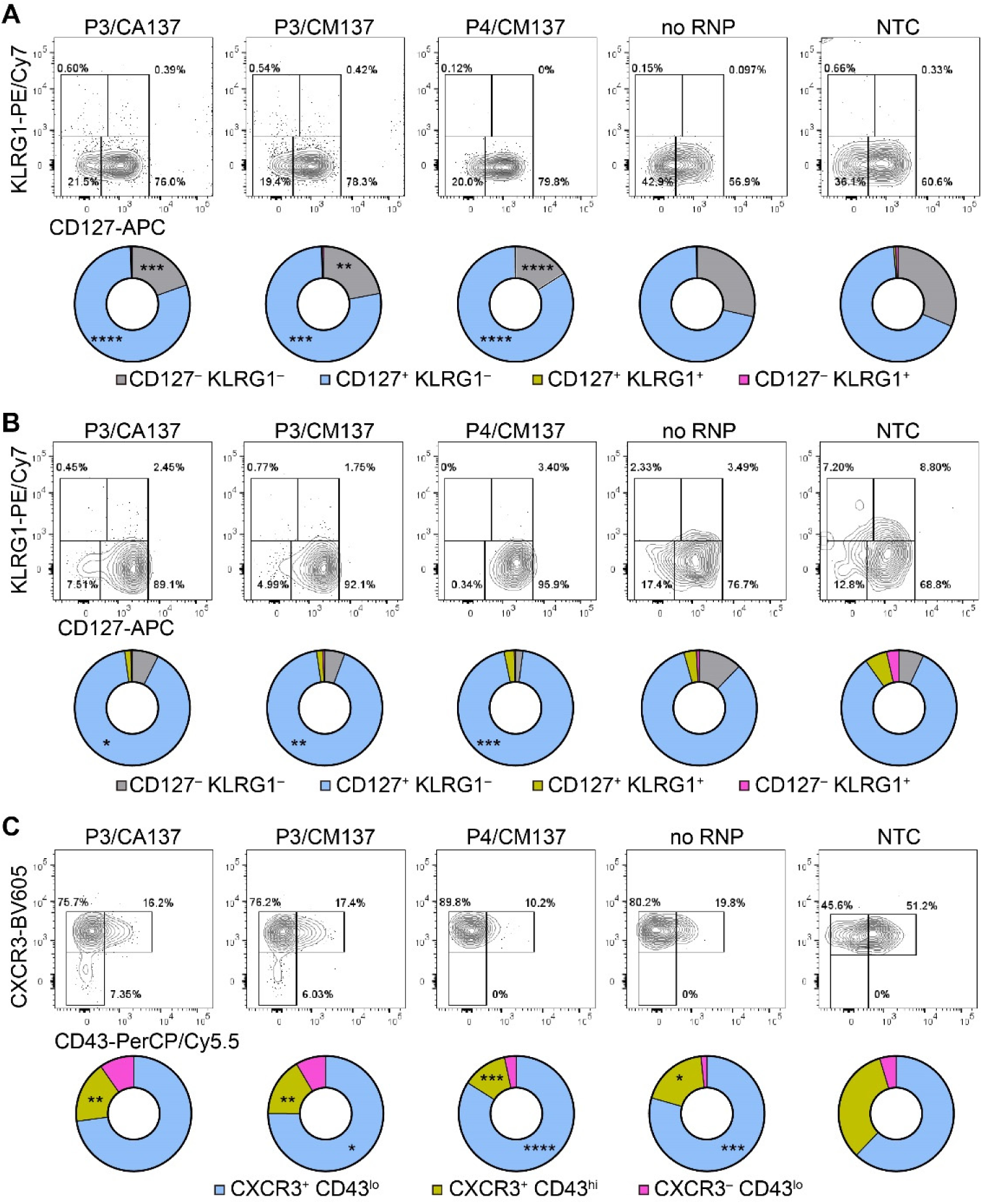
Phenotype of *in vitro*-activated OT-I cells in popLN after adoptive transfer into HSV-OVA-infected hosts. Nucleofection of OT-I cells and viral infection were performed as in **Figure 2**. (**A, B**) Expression of CD127 and KLRG1 on OT-I cells in popLN on days 7 (**A**) and > 40 (**B**). Pie charts show the mean frequencies of four populations identified by these two markers. (**C**) Expression of activation-associated glycoform of CD43 and CXCR3 on OT-I cells in popLN > 40 days after infection. Pie charts show the mean frequencies of three populations identified by these two markers. Graphs show pooled data from two independent experiments with n = 9 or 15 for nucleofected cells or non-nucleofected control, respectively. Flow cytometric plots are gated on viable OT-I cells identified by the expression of congenic markers and show concatenated data from one of two experiments with n = 5 per group. Congenic marker assignment was swapped in each experiment. NTC, non-nucleofected control. \**p* < 0.05, \*\**p* < 0.01, \*\*\**p* < 0.001, \*\*\*\**p* < 0.0001 by ordinary two-way ANOVA with Dunnett’s multiple comparison.

**Supplementary Figure 3.**
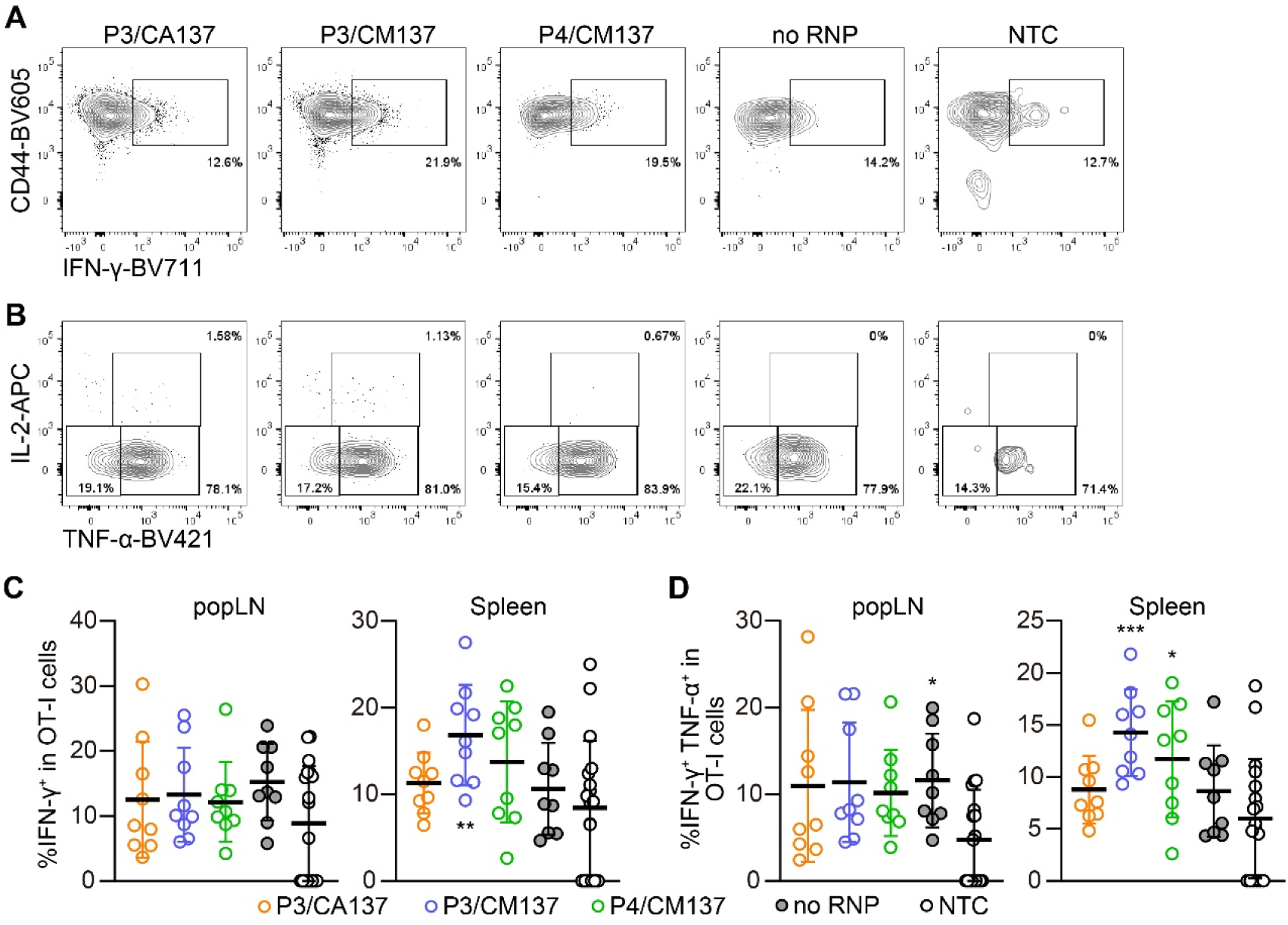
Cytokine-producing capability of *in vitro*-activated OT-I cells 3 days after adoptive transfer into HSV-OVA-infected hosts. Nucleofection of OT-I cells and viral infection were performed as in **Figure 2**. (**A, B**) Representative flow cytometric plots of IFN-γ expression in viable OT-I cells (**A**) and TNF-α and IL-2 expression among IFN-γ^+^ OT-I cells (**B**) after 5 hr restimulation with 1 µM OVA_257-264_ (SIINFEKL) peptide in the presence of brefeldin A. Flow cytometric plots show concatenated data from one of two experiments with n = 5. (**C, D**) Summary of the frequency of IFN-γ^+^ (**C**) and IFN-γ^+^ TNF-α^+^ (**C, F**) cells among viable OT-I cells. Graphs show pooled data from two independent experiments with n = 9 or 15 for nucleofected cells or non-nucleofected control, respectively. Congenic marker assignment was swapped in each experiment. NTC, non-nucleofected control. \**p* < 0.05, \*\**p* < 0.01, \*\*\**p* < 0.001 by ordinary one-way ANOVA test with Dunnett’s multiple comparison.

**Supplementary Figure 4.**
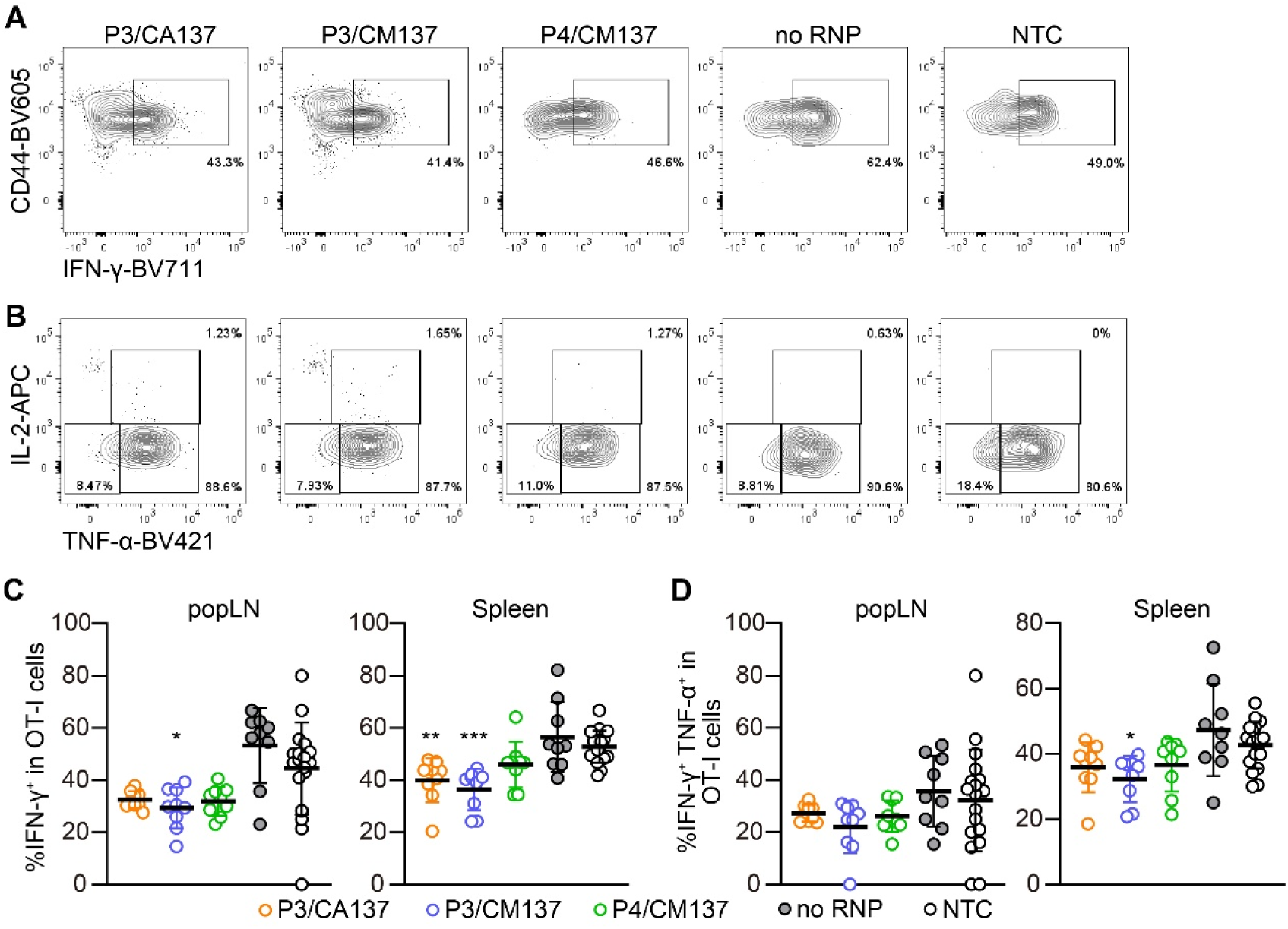
Cytokine-producing capability of *in vitro*-activated OT-I cells >37 days after adoptive transfer into HSV-OVA-infected hosts. Nucleofection of OT-I cells and viral infection were performed as in **Figure 2**. (**A, B**) Representative flow cytometric plots of IFN-γ expression in viable OT-I cells (**A**) and TNF-α and IL-2 expression among IFN-γ^+^ OT-I cells (**B**) after 5 hr restimulation with 1 µM OVA_257-264_ (SIINFEKL) peptide in the presence of brefeldin A. Flow cytometric plots show concatenated data from one of two experiments with n = 5. (**C, D**) Summary of the frequency of IFN-γ^+^ (**C**) and IFN-γ^+^ TNF-α^+^ (**C, F**) cells among viable OT-I cells. Graphs show pooled data from two independent experiments with n = 9 or 15 for nucleofected cells or non-nucleofected control, respectively. Congenic marker assignment was swapped in each experiment. NTC, non-nucleofected control. \**p* < 0.05, \*\**p* < 0.01, \*\*\**p* < 0.001 by ordinary one-way ANOVA test with Dunnett’s multiple comparison.

