## Supplemental Figures 1-4 for "Naïve and *in vitro*-activated primary mouse CD8^+^ T cells retain *in vivo* immune responsiveness after electroporation-based CRISPR/Cas9 genetic engineering"

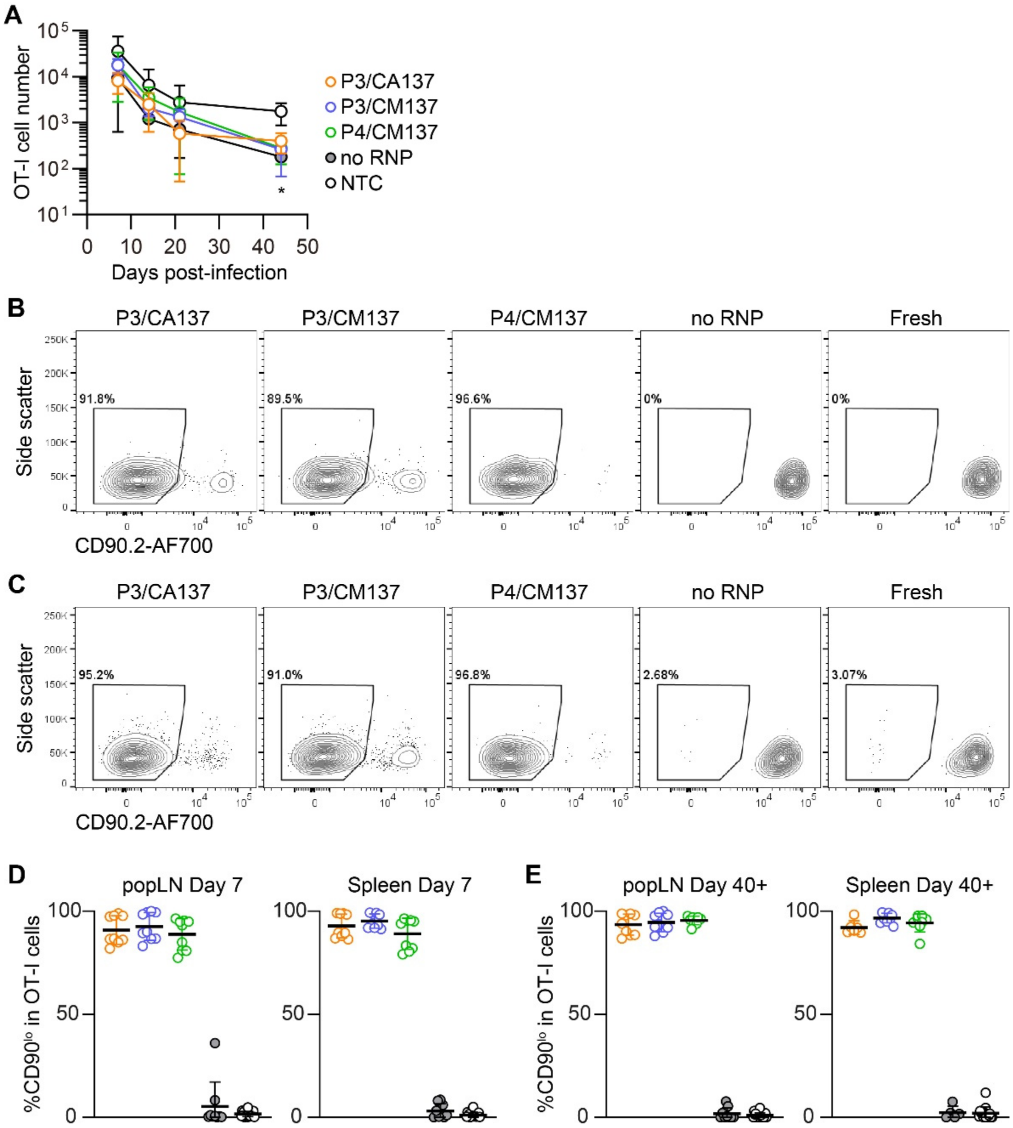


**Supplementary Figure 1.** Kinetics of the number and CD90 expression of *in vitro*-activated OT-I cells after adoptive transfer into HSV-OVA-infected hosts. Nucleofection of OT-I cells and viral infection were performed as in **Figure 2**. (**A**) Kinetics of OT-I cell number per 1 mL blood. Statistical significance of differences between each of nucleofected group and control was analyzed by ordinary two-way ANOVA with Dunnett's multiple comparison. *$p < 0.0001$. (**B–E**) Frequency of CD90$^{lo}$ cells among OT-I cells in popLN and spleen on days 7 (**B, D**) and > 40 (**C, E**). Congenic marker assignment was swapped in each experiment. Data are pooled from two independent experiments with n = 6–9 or n = 12–15 for nucleofected cells or non-nucleofected control, respectively.

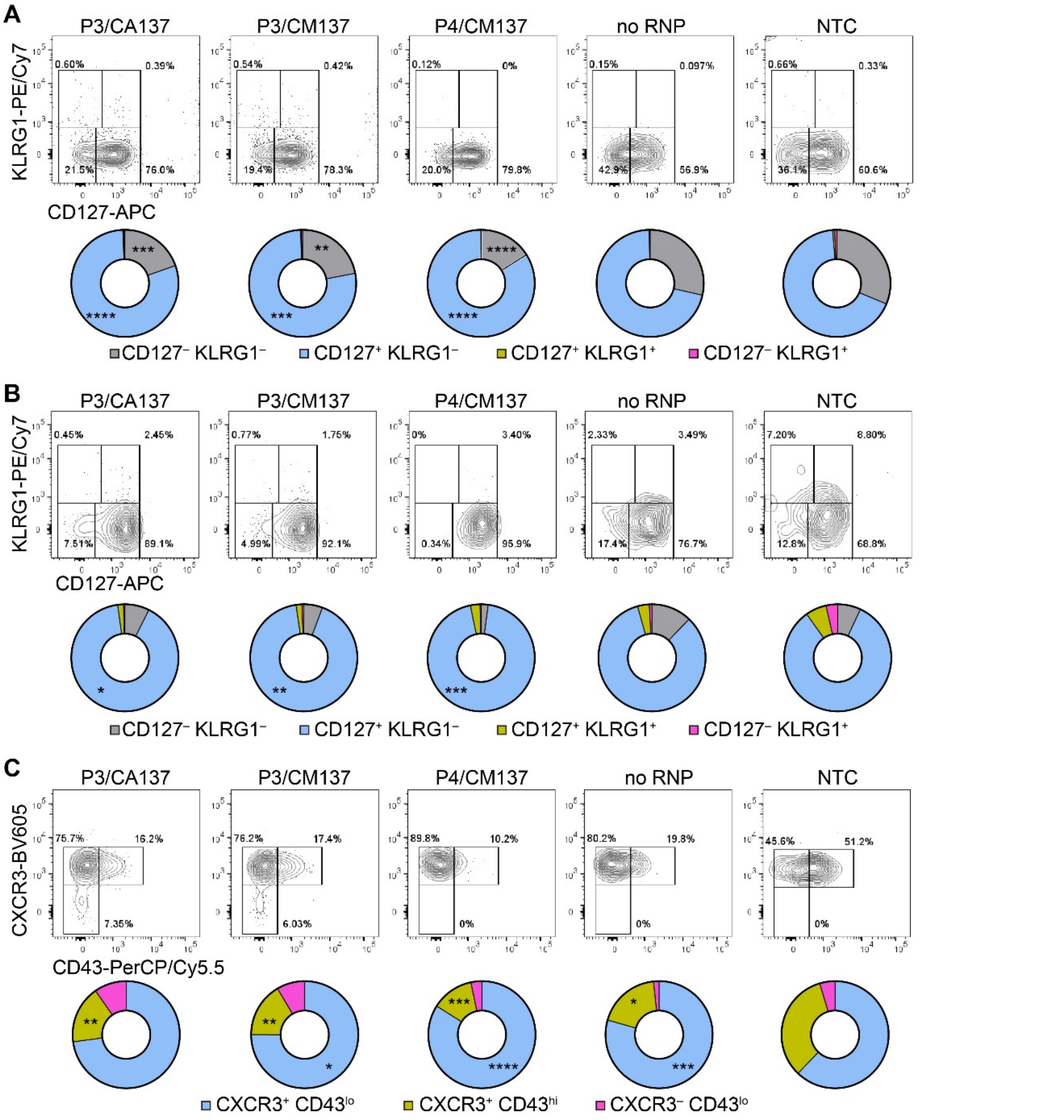


**Supplementary Figure 2.** Phenotype of *in vitro*-activated OT-I cells in popLN after adoptive transfer into HSV-OVA-infected hosts. Nucleofection of OT-I cells and viral infection were performed as in **Figure 2**. (**A, B**) Expression of CD127 and KLRG1 on OT-I cells in popLN on days 7 (**A**) and > 40 (**B**). Pie charts show the mean frequencies of four populations identified by these two markers. (**C**) Expression of activation-associated glycoform of CD43 and CXCR3 on OT-I cells in popLN > 40 days after infection. Pie charts show the mean frequencies of three populations identified by these two markers. Graphs show pooled data from two independent experiments with n = 9 or 15 for nucleofected cells or non-nucleofected control, respectively. Flow cytometric plots are gated on viable OT-I cells identified by the expression of congenic markers and show concatenated data from one of two experiments with n = 5 per group. Congenic marker assignment was swapped in each experiment. NTC, non-nucleofected control. **p* < 0.05, ***p* < 0.01, ****p* < 0.001, *****p* < 0.0001 by ordinary two-way ANOVA with Dunnett's multiple comparison.

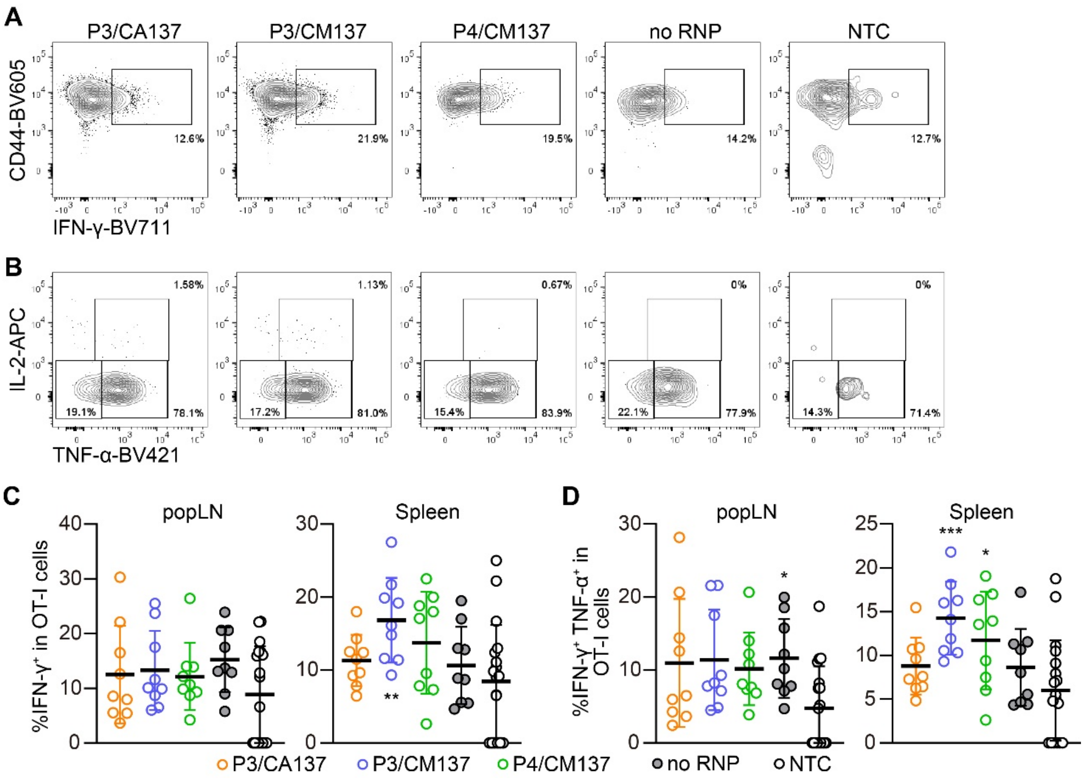


**Supplementary Figure 3.** Cytokine-producing capability of *in vitro*-activated OT-I cells 3 days after adoptive transfer into HSV-OVA-infected hosts. Nucleofection of OT-I cells and viral infection were performed as in **Figure 2**. (**A, B**) Representative flow cytometric plots of IFN-γ expression in viable OT-I cells (**A**) and TNF-α and IL-2 expression among IFN-$\gamma^+$ OT-I cells (**B**) after 5 hr restimulation with 1 μM $OVA_{257-264}$ (SIINFEKL) peptide in the presence of brefeldin A. Flow cytometric plots show concatenated data from one of two experiments with n = 5. (**C, D**) Summary of the frequency of IFN-$\gamma^+$ (**C**) and IFN-$\gamma^+$ TNF-$\alpha^+$ (**C, F**) cells among viable OT-I cells. Graphs show pooled data from two independent experiments with n = 9 or 15 for nucleofected cells or non-nucleofected control, respectively. Congenic marker assignment was swapped in each experiment. NTC, non-nucleofected control. *$p < 0.05$, **$p < 0.01$, ***$p < 0.001$ by ordinary one-way ANOVA test with Dunnett's multiple comparison.

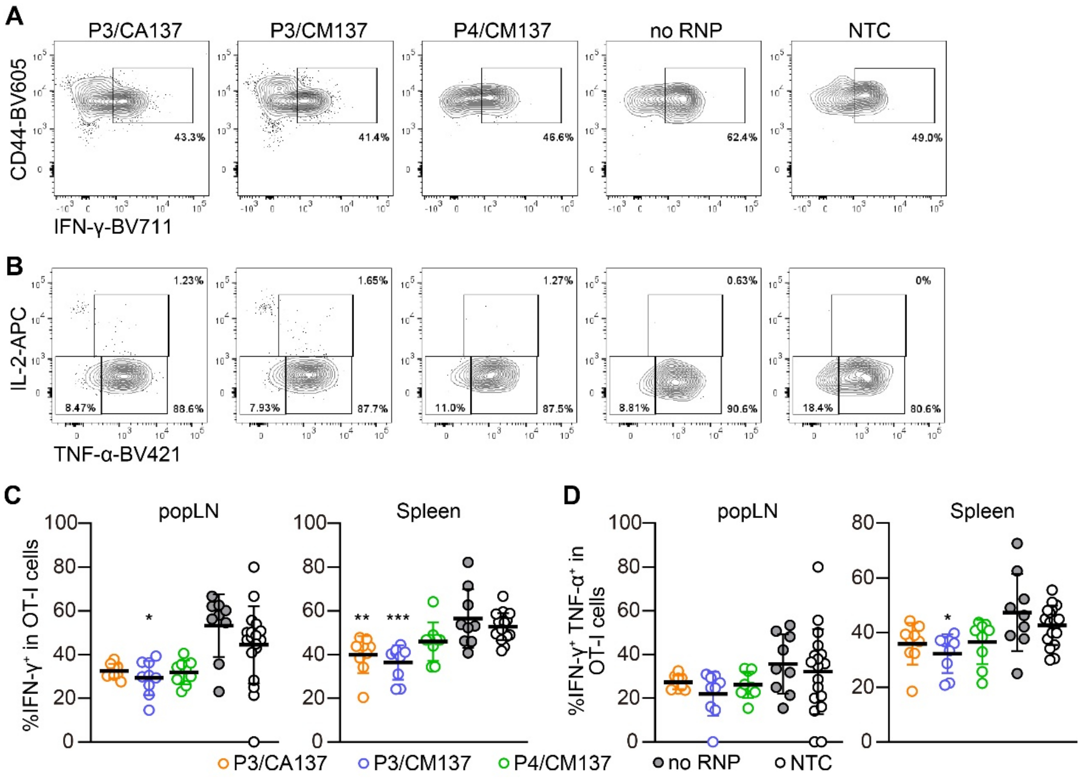


**Supplementary Figure 4.** Cytokine-producing capability of *in vitro*-activated OT-I cells >37 days after adoptive transfer into HSV-OVA-infected hosts. Nucleofection of OT-I cells and viral infection were performed as in **Figure 2**. (**A, B**) Representative flow cytometric plots of IFN-γ expression in viable OT-I cells (**A**) and TNF-α and IL-2 expression among IFN-$\gamma^+$ OT-I cells (**B**) after 5 hr restimulation with 1 μM $OVA_{257\text{-}264}$ (SIINFEKL) peptide in the presence of brefeldin A. Flow cytometric plots show concatenated data from one of two experiments with n = 5. (**C, D**) Summary of the frequency of IFN-$\gamma^+$ (**C**) and IFN-$\gamma^+$ TNF-$\alpha^+$ (**C, F**) cells among viable OT-I cells. Graphs show pooled data from two independent experiments with n = 9 or 15 for nucleofected cells or non-nucleofected control, respectively. Congenic marker assignment was swapped in each experiment. NTC, non-nucleofected control. **p* < 0.05, ***p* < 0.01, ****p* < 0.001 by ordinary one-way ANOVA test with Dunnett's multiple comparison.
